# Molecular Basis of Far-red Sensing in Cyanobacteriochrome

**DOI:** 10.1101/2020.06.02.130930

**Authors:** Sepalika Bandara, Nathan Rockwell, Xiaoli Zeng, Zhong Ren, Cong Wang, Heewhan Shin, Shelley S. Martin, Marcus V. Moreno, J. Clark Lagarias, Xiaojing Yang

## Abstract

Cyanobacteriochromes are small, panchromatic photoreceptors in the phytochrome superfamily that regulate diverse light-mediated adaptive processes in cyanobacteria. The molecular basis of far-red (FR) light perception by cyanobacteriochromes is currently unknown. Here we report the crystal structure of a far-red-sensing cyanobacteriochrome from *Anabaena cylindrica* PCC 7122, which exhibits a reversible far-red/orange photocycle. The 2.7 Å structure of its FR-absorbing dark state, determined by room temperature serial crystallography and cryo-crystallography, reveals an *all-Z,syn* configuration of its bound linear tetrapyrrole (bilin) chromophore that is less extended than the bilin chromophores of all known phytochromes. Based on structural comparisons with other bilin-binding proteins and extensive spectral analyses on mutants, we identify key protein-chromophore interactions that enable far-red sensing in bilin-binding proteins. We propose that FR-CBCRs employ two distinct tuning mechanisms, which work together to produce a large batho-chromatic shift. Findings of this work have important implications for development and improvement of photoproteins with far-red absorption and fluorescence.

**Significance Statement:** Phytochromes are well known far-red-light sensors found in plants that trigger adaptive responses to facilitate competition for light capture with neighboring plants. Red- and far-red-sensing are critical to cyanobacteria living in the far-red-enriched shade of plants. Here we report the crystal structure of a far-red-sensing cyanobacteriochrome, a distant cyanobacterial relative of phytochrome. These studies shed insight into the poorly understood molecular basis of far-red-sensing by phytobilin-based photoreceptors. Owing to the deep tissue penetration of far-red light, far-red-sensing photoreceptors offer promising protein scaffolds for developing gene-based photoswitches, optoacoustic contrast agents and fluorescent probes for *in situ* imaging and optogenetic applications.

## Introduction

Cyanobacteria have developed elaborate, spectrally tuned photoreceptors and light-harvesting systems for adaptation and survival in a wide range of ecological niches (1–5). Many photoreceptor systems are modular components of much larger signaling proteins that integrate different sensor and effectors modules in a single protein molecule and interface with diverse signal transduction pathways. Photoreceptors in the phytochrome superfamily utilize a specific lineage of the GAF (c**G**MP phosphodiesterase, **A**denylyl cyclase and **F**hlA) signaling module to bind a thioether-linked linear tetrapyrrole (bilin) chromophore for light perception via *15,16*-photoisomerization (6–11). Such photoreceptors play critical roles in plant development as well as in regulating cyanobacterial phototaxis, development and light harvesting (2, 3, 12–17). Protein structural changes following the primary photochemical event then alter the downstream enzymatic activities and/or protein-protein interactions via an interdomain allosteric mechanism (18).

Phytochromes possess a tripartite photosensory module consisting of three N-terminal domains (PAS, GAF and PHY), known as the photosensory module, in which the PAS and GAF domains are tethered via a “figure-of-eight knot” (14, 19, 20). In typical phytochromes, the bilin chromophore embedded in the GAF domain adopts a *5-Z,syn, 10-Z,syn, 15-Z,anti* configuration in the dark-adapted state. Light absorption triggers photoisomerization of the 15,16–double bond to generate a *15E,anti* photoproduct, which typically absorbs far-red light (9, 14, 21). An long extension from the adjacent PHY domain is responsible for stabilizing the far-red-absorbing Pfr state (14, 20). In cyanobacteria, the phytochrome superfamily has diversified to yield a large family of more streamlined sensors, designated cyanobacteriochromes (CBCRs) (2, 4, 22–26). Unlike canonical phytochromes, CBCR photosensory modules consist of one or more stand-alone GAF domains that are sufficient for covalent attachment of bilin and photoconversion. These small CBCR domains have also been used as light-sensing modules in a variety of synthetic biology applications (27–32). In contrast to the typical red/far-red phytochromes, CBCRs are able to sense all colors of light from near UV to far-red utilizing a common phycocyanobilin (PCB) chromophore precursor (22–24, 26).

The remarkable spectral diversity of CBCRs arises from extensive molecular evolution of the GAF domain scaffold. Many CBCRs leverage two thioether linkages to sense blue, violet or near-UV light (8, 22, 23, 25, 33–35). Such ‘two-Cys’ CBCRs possess an additional thioether linkage to the C10 methine bridge of the bilin that splits the chromophore in half, significantly shortening the conjugated π system. Rupture of this covalent bond can occur upon 15Z/15E photoisomerization, which restores bilin conjugation across C10 to generate a photo-state absorbing at wavelengths from teal to red (8, 33, 36, 37). Dual cysteine CBCRs have evolved multiple times, yielding a wide range of photocycles with violet, blue, teal, green, orange and red states (22).

Other CBCRs use different tuning mechanisms. Red/green CBCRs such as AnPixJg2 and NpR6012g4 have a red-absorbing dark state that photoconverts to a green-absorbing lit state. In this subfamily, the molecular mechanism responsible for photoproduct tuning relies on trapping the *15E* bilin in a twisted geometry that results in blue-shifted absorption (10, 11). In contrast, green/red CBCRs have a reversed photocycle: the green-absorbing *15Z* dark state photoconverts to yield a red-absorbing *15E* photoproduct. This subfamily uses a protochromic mechanism, whereby photoconversion triggers a proton transfer to the chromophore and induces a spectral red shift (2, 38).

Until recently, the light sensing range of CBCRs appeared limited to the visible spectrum thereby implicating phytochromes to be exclusively responsible for FR sensing. Indeed, far-red-dependent remodeling of the photosynthetic apparatus in multiple cyanobacterial species is mediated by the phytochrome RfpA (3, 39). The discovery of two lineages of CBCRs with FR-absorbing dark states (FR-CBCRs) was thus surprising (40). Upon absorbing far-red light, such FR-CBCRs convert to either an orange- or red-absorbing photoproduct state. FR-CBCRs have evolved from green/red CBCRs as part of a Greater Green/Red (GGR) lineage, diversification comparable to that seen in the XRG (extended red/green) lineage (35, 40, 41). Owing to their small size and spectral overlap with the therapeutic window of optimum tissue penetrance (700-800 nm) (42–46), FR-CBCRs provide a tantalizing scaffold for development of FR-responsive optogenetic reagents for biomedical research and clinical applications (45, 47–50).

To understand the molecular basis of far-red spectral tuning, we determined the crystal structures of the FR-absorbing dark state of a representative FR/O CBCR at both ambient and cryogenic temperatures. These structures revealed an *all-Z,syn* configuration of its bilin chromophore that is different from those found in all other CBCR or phytochrome structures. Based on these crystallographic results together with spectral analysis of mutants and related FR-CBCRs, comparisons with other bilin-binding proteins, we identify key protein-chromophore interactions, which suggest two distinct tuning mechanisms simultaneously at work for far-red light detection in FR-CBCRs.

## Results and Discussion

### Crystal structure of 2551g3 reveals an unusual all-Z,syn chromophore

We have determined the structure of a representative FR-CBCR Anacy_2551g3 (hereafter, 2551g3) (40). 2551g3 is the third GAF domain of a multi-domain sensor histidine kinase encoded by locus Anacy_2551 from *Anabaena cylindrica* PCC 7122 (SI Fig. S1A). This FR-CBCR as a truncated GAF-only construct covalently incorporates the phycocyanobilin (PCB) chromophore when it is co-expressed with a PCB-producing plasmid in *E. coli* (40, 51, 52) and exhibits reversible photoconversion between a FR-absorbing *15Z* dark state (Pfr; λmax 728 nm) and an orange-absorbing *15E* lit state (Po; λmax 588 nm; SI Fig. S2A). The crystal structure of 2551g3 in the Pfr dark state was determined at both cryogenic and room temperatures (Fig. 1; SI Materials and Methods). Retention of the Pfr state in the crystal was confirmed by single-crystal absorption spectroscopy and by the characteristic green color of 2551g3 crystals (SI Fig. S2B). Monochromatic datasets collected at 100 K were used to obtain an initial model for 2551g3 at 3.1 Å resolution by the molecular replacement method (PHENIX) using the crystal structure of the two-Cys CBCR TePixJ (PDB: 4GLQ) (8) as a search model. To improve the resolution and map quality, we employed an *in situ* room-temperature (RT) data collection method developed by our laboratory (53), which yielded a complete Laue dataset at 2.7 Å resolution from >800 crystals of 2551g3. Our serial RT crystallography strategy not only alleviates X-ray radiation damage but also evades crystal freezing, particularly beneficial given a high solvent content of ∼70% in 2551g3 crystals. The resulting high-quality electron density map allowed us to build a complete 2551g3 structure with no gaps in the protein backbone. Compared to the cryo-crystallography data, this room temperature (RT) structure shows excellent statistics both in the real space and reciprocal space (SI Table S1) with well-resolved electron density for the bilin chromophore (Fig. 2A). Unless mentioned otherwise, the RT structure of 2551g3 is used for depiction and discussion in this work.

**Figure 1.**
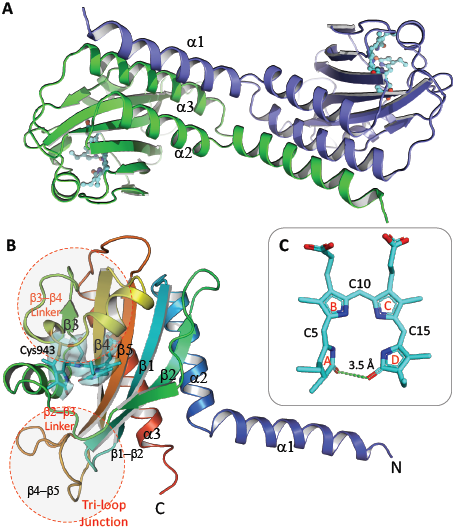
Crystal structure of far-red CBCR 2551g3 in the Pfr state. **(A)** In a “handshake” dimer structure of 2551g3, the GAF-α1 helix of one protein molecule (green) extends out to form a three-helix bundle with the GAF-α2 and GAF-α3 helices from the partner molecule (blue). **(B)** Phycocyanobilin (PCB; cyan) is located in a cleft sandwiched by the β3−β4 and β2−β3 linkers. The electron densities of the 2Fo-Fc map (rendered in transparent cyan surface at the 2.2σ contour) are shown for PCB and the Cys943 anchor. Two structural features unique to the FR-CBCR family, a disjointed helix at the α-face and tri-loop junction at the β-face, are highlighted in red circles. **(C)** The PCB chromophore adopts an unusual *all-Z,syn* conformation.

**Figure 2.**
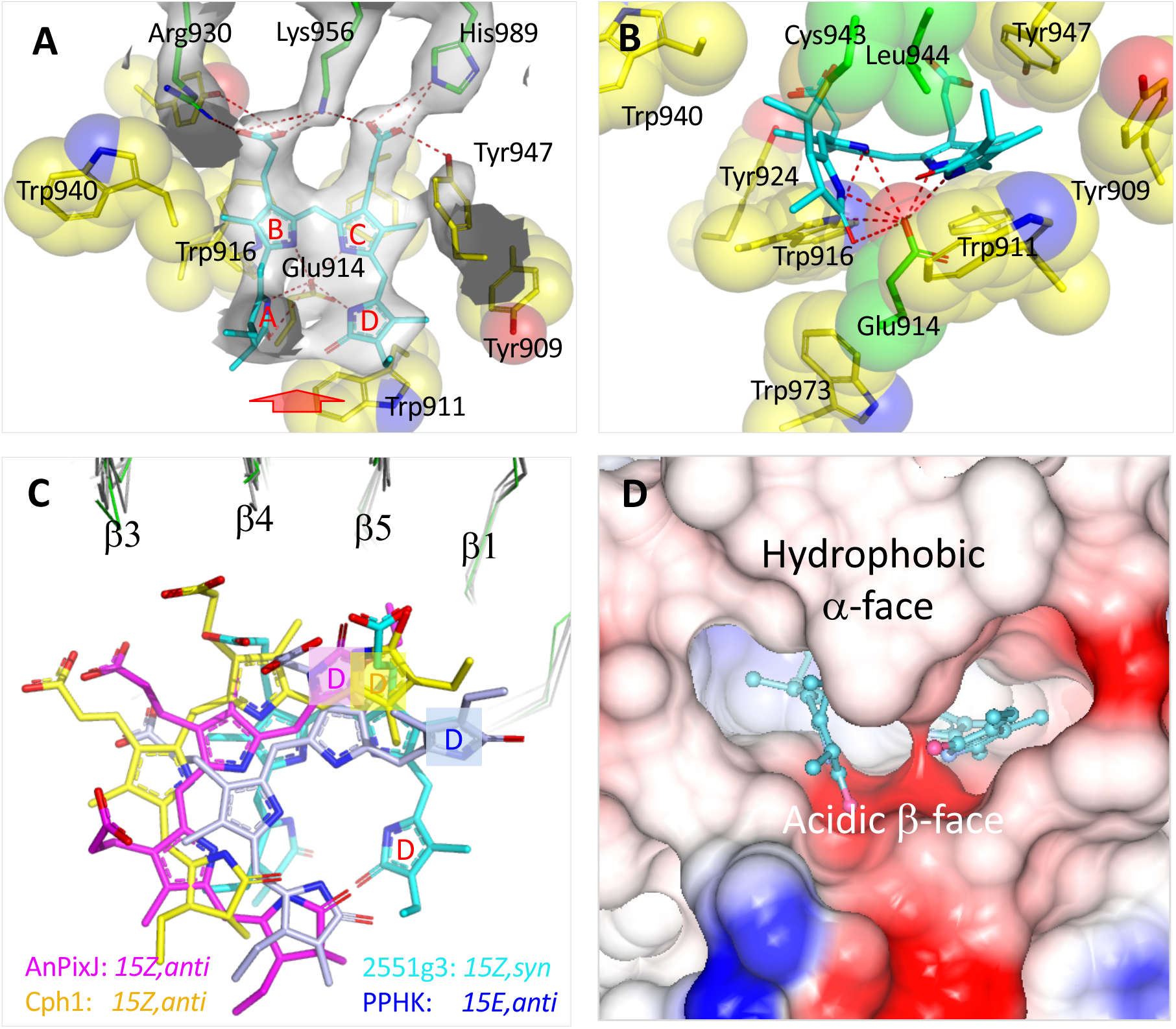
The chromophore binding pocket in the Pfr structure of 2551g3. **(A)** The 2*F*o-*F*c map (contoured at 1.7 σ; rendered in gray transparent surface) shows close interactions between the propionates of PCB (cyan) and its protein anchors (Arg930/Lys956, His989/Tyr947; in green sticks). **(B)** Viewed from the red arrow in panel A, Leu944 is located next to the Cys943 anchor at the α-face, whereas both rings A and D point towards the β-face. Red dashed lines highlight the hydrogen bonds between PCB and the acidic side chain of the hallmark residue Glu914. Yellow spheres mark the surrounding aromatic residues. **(C**) PCB in the Pfr structure of 2551g3 (cyan) assumes a distinct disposition relative to other phytochrome systems: canonical phytochrome Cph1 (2VEA: yellow), red/green CBCRs AnPixJ (3W2Z: magenta) and PPHK (6OAP: light blue). All structures are aligned based on the GAF protein framework (shown in thin wires). **(D**) The electrostatic surface in the PCB cavity is highly asymmetric between the α− and β−faces.

The Pfr structure of 2551g3 in the P4_2_22 space group contains one protein molecule in an asymmetric unit (Fig. 1). The core domain features a typical GAF fold with a central β-sheet consisting of five β strands in a topological order of β2−β1−β5−β4−β3 (Fig. 1B). However, the GAF-α1 helix clearly disengages from its own protomer and bundles with the GAF-α2 and GAF-α3 helices from a symmetry-related protomer, forming a “handshake” crystallographic dimer (Fig. 1A). 2551g3 and related CBCRs are characterized by a large insertion between the β4 and β5 strands (aa. 964 -980) (40), in contrast to a tight turn typically found in other CBCRs and prokaryotic phytochromes (SI Fig. S3A). This large insertion folds back towards the core and directly interacts with the β2-β3 linker and the β1−β2 loop, constituting a “tri-loop junction” structure located on the β-face of the chromophore (54) (Fig. 1B). On the α-face of PCB, two short helices in the β3-β4 linker are connected by a sharp turn containing Cys943, the conserved Cys to which the chromophore is covalently attached. This region is thus structurally distinct from phytochromes and CBCRs that place the Cys anchor at the start of a continuous three-turn helix (SI Fig. S3B).

The chromophore pocket of 2551g3 is surrounded by aromatic residues conserved in other CBCRs (Fig. 2A and B, yellow). The β3−β4 and β2−β3 linkers, which are located respectively at the α and β faces of the bilin, form a cleft that shields the chromophore (Fig. 1B). The PCB chromophore, which is covalently attached to Cys943 via the ring A ethylene group, adopts a compact *all-Z,syn* conformation. In this cyclic conformation, rings A and D are tilted towards the β-face with respect to rings B and C (Fig. 1B, 1C). Their ring planes are nearly perpendicular to each other with two carbonyl oxygen atoms separated by ∼3.5 Å (Fig. 1C). This conformation results in unique dihedral angles about the C5 and C15 methine bridges compared to 156 bilin-binding protein structures in the Protein Data Bank (SI Fig. S4). The 2551g3 chromophore also adopts a much more distorted conformation than the *all-Z,syn* bilins bound to proteins such as ferredoxin-dependent bilin reductase (FDBR) PcyA and bilin lyase CpcT (SI Fig. S5) (55, 56). The covalently attached bilins in the previously characterized phytochromes, CBCRs, and light-harvesting phycobiliproteins adopt extended *15-Z,anti* or *15-E,anti* configurations (7, 8, 10, 11, 14, 19, 20, 57, 58), which would cause major steric clashes with the chromophore binding pocket in 2551g3 (SI Fig. S6), despite the *5-Z,syn* configuration in all cases. These clashes are in part reconciled in 2551g3 as the chromophore in 2551g3 rotates clockwise in the approximate plane of the bilin ring system when viewed from the chromophore α-face. To our best knowledge, ring D of the *all-Z,syn* bilin in 2551g3 assumes the most extreme clockwise orientation relative to the GAF framework among known phytochromes and CBCRs (Fig. 2C).

### The all-Z,syn bilin engages unique protein-chromophore interactions

The 2551g3 structure reveals several unique protein-chromophore interactions in FR-CBCRs and related CBCRs (SI Fig. S1B). *First*, Leu944 next to the anchor Cys943 replaces a histidine residue highly conserved among the phytochrome superfamily (Fig. 2B) (40). The hydrophobic Leu944 approaches the bilin from a disjointed α-facial helix, resulting in tilting of rings A and D towards the β-face (Fig. 2B, 2D). *Second*, the hallmark residue Glu914 at the β-face exhibits hydrogen bonds with all four pyrrole nitrogen atoms, yet evidently closer to rings C and D according to the electron density map (Fig. 2A). The His→Leu substitution at the α-face and the acidic β-facial patch constitute a highly asymmetric protein environment for the pyrrole rings, where both bilin lactam groups from ring A and D point to the acidic β-face (Fig. 2D). *Third*, aromatic residues conserved among FR-CBCRs CBCRs play a steric role in generating a snug protein pocket of 2551g3 where all bilin rings are constrained in the *all-Z,syn* conformation (40) (Fig. 2A-B). Ring A is repelled toward the β-face by Leu944, forming hydrogen bonds with Glu914, Trp916 and the protein backbone atoms. Rings B and C are laterally clamped between Ile939/Trp940 and Tyr924 while engaging close steric interactions with Tyr916, Leu944, Phe884 and Tyr947 at both α- and β-faces. Ring D is stacked against Trp911, a position often occupied by an aromatic residue in FR-CBCRs and related CBCRs (Fig. S1B). It is likely that such steric factors preclude binding of any bilins in the extended *15,anti* conformation (Fig. S6).

In addition to steric constraints, the bilin propionates engage ionic interactions with basic residues deep within the GAF β sheet. Specifically, Lys956 and Arg930 form salt bridges with the ring B propionate, while His989 and Tyr947 are well positioned to interact with the ring C propionate (Fig. 2A). The electron densities associated with the ringC-His989 and ringC-Lys956 interactions are visible in the *2Fo-Fc* map of 2551g3 contoured at 3σ (Fig. 2A). In phytochromes and CBCRs, a conserved His residue at the equivalent position of His989 is important for stabilizing the ring D in the *15Z,anti* conformation (7, 8, 10, 19, 20), while interacting with the ring C propionate in the *15E,anti* conformation (14, 21) (SI Fig. S6). This interaction between His989 and the C-ring propionate is thus characteristic to FR-CBCRs, where it plays a role in stabilizing the compact *all-Z,syn* chromophore.

### Critical residues for FR sensing identified by mutagenesis

To identify key protein-chromophore interactions for FR absorption, we carried out site directed mutagenesis on 2551g3 and a closely related FR-CBCR Anacy_4718g3 (hereafter, 4718g3; SI Fig. S1B) (40). A total of 34 variant proteins with substitutions occurring at 25 different positions were evaluated. 20 variants exhibited a range of severe phenotypes including loss of bilin binding, loss of the far-red state, and major changes in photoproduct tuning (SI Fig. S7). Other variants only have minor effects on peak wavelength and/or photoproduct lineshape. Among nine variants that failed to bind bilin are single Leu substitutions at the positions of His989, Lys956, Arg930 and Asp938, which anchor the bilin propionates (Fig. 2A, SI Fig. S8). The L944H variant of 2551g3, which restores the α-facial Histidine residue typically found at the equivalent position in phytochromes and CBCRs, did not bind bilin. In contrast, the L1383N variant of 4718g3 at the equivalent position binds PCB, but at a reduced level with a blue-shifted peak absorption of 590 nm (SI Fig. S7). It is possible that the slightly bulkier side chain of His prevents proper bilin attachment.

Only three variants disrupted far-red absorption. In addition to the L1383N of 4718g3, the E914D variant of 2551g3 also exhibited reduced chromophorylation with a red-absorbing dark state below 700 nm, and photoconversion to the photoproduct is also limited (SI Fig. S7, S8). However, the equivalent E1353Q and E1353D variants of 4718g3 did not even bind PCB. Substitution of the adjacent Asp913 for Leu resulted in a larger blue shift, again with reduced chromophorylation, although this residue is not strictly conserved among FR-CBCRs. In the 2551g3 structure, Asp913 forms a salt bridge to Arg972 at the tri-loop junction. The R972A variant (R1410A of 4718g3), however, retain both FR absorption and photoconversion (SI Fig. S7, S8), so is the E912N variant (D1351N of 4718g3). These results suggest that the FR-absorbing phenotype is only affected by mutations at the positions of Leu944 and Glu914, which are the hallmark residues in FR-CBCRs. In the 2551g3 structure, they are positioned to directly influence the protonation state of the bilin pyrrole rings (Fig. 2B).

Eight substitutions resulted in red-shifted photoproducts (SI Fig. S7). These include single mutations at several bulky residues specific to FR-CBCRs. Some variants exhibited far-red/red photocycles while others yielded both orange- and red-absorbing photoproducts upon photoconversion (SI Fig. S7). Substitutions at Trp911 had only minor effects on spectral tuning in 2551g3, while the corresponding W1350A and W1350F variants of 4718g3 exhibited normal far-red-absorbing states with red-shifted light states, as did W911F of 2551g3. Such far-red/red photocycles were also seen in W940L of 2551g3, W1379L and I1378P of 4718g3 as well as in the S917P/Q918P double mutant. On the other hand, mixed *15E* photoproducts were observed with substitutions at Tyr947 and the equivalent Tyr1386 of 4718g3 (SI Fig. S7C). We note that the wild-type 4718g3 exhibited a small amount of a red-absorbing photoproduct species as a shoulder (SI Fig. S7B). These data supports that FR-CBCRs have a continuum of possible photoproducts.

With few exceptions, the Pfr dark state is largely retained in the Trp variants (SI Fig. S7), suggesting that individual Trp residues are not directly involved the far-red tuning. Rather, they play a role in formation of the Po photoproduct state, which may assume a twisted *15E* conformation following the 15,16-photoisomerization. With steric constraints removed by mutagenesis at these Trp positions, the *15E* chromophore would be allowed to adopt a more relaxed conformation rendering a red-shifted photoproduct (40).

### FR-CBCRs have a protonated far-red-absorbing state

The high sequence similarity between FR-CBCRs and green/red CBCRs (Fig. S1B) suggest that FR-CBCRs have a protochromic photocycle as observed in green/red CBCRs RcaE and CcaS where the chromophore is deprotonated in the green-absorbing *15Z* photostate but becomes protonated in the *15E* photoproduct (2). We therefore carried pH titration experiments to examine the protonation state of FR-CBCRs. The Pfr state of 2551g3 is quite stable over a wide range of pH from 5 to 10 (Fig. 3C). At pH > 10, 2551g3 displayed pH-dependent conversion to the orange-absorbing state, which is fully reversible by lowering pH. To confirm this protochromic behavior, we examined two additional FR-CBCRs, Syn7502_01757 (hereafter, 01757) and WP_046814686 (hereafter, 0468g3), which were identified by an updated phylogenetic analysis (SI Fig. S9A) Both represent early-diverging members in the FR-CBCR cluster to which 2551g3 and 4718g3 belong (40) (SI Fig. S9B). As expected, both proteins exhibited far-red *15Z* photostates. However, 01757 generated a green-absorbing photoproduct, and 0468g3 exhibited a red-absorbing photoproduct (SI Fig. S9D-E), in contrast to the orange-absorbing photoproduct as in 2551g3 and 4718g3. Photoconversion of acid-denatured samples confirmed that these proteins underwent the 15,16-photoisomerization reaction (SI Fig. S9F). Given the diversity of photoproducts manifested in the FR-CBCR clade as well as the 2551g3 mutants, we re-designate this lineage as the FR/X CBCRs (SI Fig. S7C, SI Fig. S9A).

**Figure 3.**
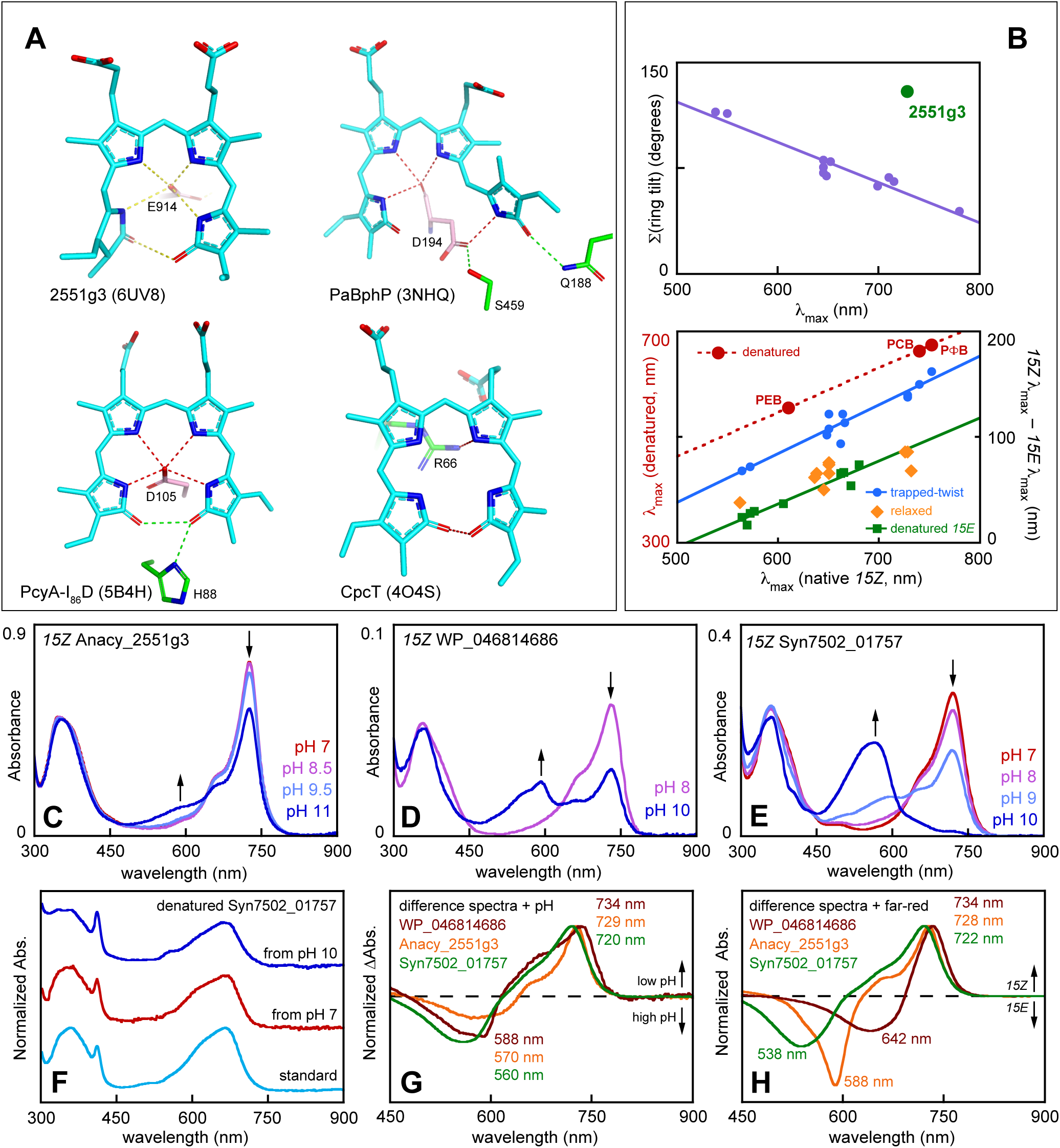
A protonated chromophore in the far-red absorbing state. **(A)** Selected protein-chromophore interactions are shown in dashed lines for the indicated structures, along with the corresponding PDB code. **(B)** *Top*, ring tilt angles were calculated using an in-house script for adjacent rings about each methine bridge. The sum of the three angles was plotted against peak wavelength for a range of XRG CBCRs and for I_86_D PcyA (purple; linear fit, r^2^ = 0.92). 2551g3 (green) is shown for comparison. **(C)** *Bottom*, correlations are shown between different spectral parameters for CBCRs, using data reported in [#@41]. Peak wavelengths are plotted for denatured and native samples of 4718g3 incorporating different bilins (large red circles, dashed curve; linear fit, r^2^ > 0.99). Other curves show trapped-twist (small blue circles; linear fit, r^2^ = 0.92), relaxed (orange diamonds), and denatured (green squares; linear fit, r^2^ = 0.93) CBCRs incorporating a range of bilins, with the blue-shift upon photoconversion plotted versus the native *15Z* peak wavelength. **(C)** Absorbance spectra for *15Z* 4718g3 are shown at pH 7 (red) and pH 10 (dark blue). **(D)** Absorbance spectra for *15Z* 0468g3 are shown at pH 8 (mauve) and pH 10 (dark blue). **(E)** Absorbance spectra for *15Z* 01757 are shown at pH 7 (red), pH 8 (mauve), pH 9 (periwinkle), and pH 10 (dark blue). **(F)** Normalized absorbance spectra are shown for samples of *15Z* 01757 at standard pH 7.8 (teal), pH 7 (red), and pH 10 (dark blue) that were then denatured. **(G)** Normalized difference spectra are shown for *15Z* FR-CBCRs upon pH change: 4718g3 (orange, calculated from panel C), 0468g3 (brick red, calculated from panel D), and 01757 (green, calculated from extrema in panel E). **(H)** Normalized photochemical difference spectra are shown for 4718g3 (orange, calculated from SI Fig. S7B, *top*), 0468g3 (brick red, calculated from SI Fig. S8E), and 01757 (green, calculated from SI Fig. 8D).

These two previously uncharacterized FR/X CBCRs indeed confirmed the presence of a protonated chromophore in the far-red-absorbing state. At pH10, 0468g3 exhibited approximately 50% loss of far-red absorption with a clear absorption rise in the yellow to orange region (Fig. 3D). 01757 exhibited nearly complete loss of the far-red band, with appearance of a well-resolved band in the green to orange region while retaining the *15Z* configuration (Fig. 3E-F). In addition, difference spectra induced by pH change are remarkably similar among FR-CBCRs, although they have different photoproducts and vary in the extent of the pH-induced transition (Fig. 3G-H). These results suggest that 2551g3 and related FR-CBCRs have a protonated bilin in the far-red state.

### Far-red-absorbing mechanisms in FR-CBCRs

To identify the molecular basis for far-red sensing, we compared the 2551g3 structure with those of other bilin-binding proteins that also absorb far-red light, in particular, bacteriophytochrome PaBphP and an I86D variant of ferredoxin-dependent bilin reductase PcyA (14, 59) (Fig. 3). Both bind biliverdin (BV) instead of PCB. We note that all three far-red structures feature a hydrogen bond between an Asp or Glu residue (corresponding to Glu914 of 2551g3) and the D-ring NH moiety. PcyA places the conserved Asp105 in close proximity to all four NH moieties of BV in an *all-Z,syn* configuration (55, 59). In PaBphP, the carboxylate side chain of Asp194 from the PxSDIP motif directly interacts with the D-ring of BV in the *5-Z,syn 10-Z,syn 15-E,anti* conformation. Both PcyA and 2551g3 have an *all-Z,syn* bilin. While the BV chromophore in PcyA-I86D largely follows the general trend of bilin twist vs. peak wavelength demonstrated in bilin-binding proteins, the twisted PCB chromophore of 2551g3 does not follow the same trend (Fig. 3B). However, we note a different correlation between the Q-band peak wavelength vs. pKa of the residue interacting with the D-ring NH group. For example, the Pr state is observed in phytochromes and red/green CBCRs where the D-ring NH moiety of the *15Za* chromophore forms a hydrogen bond with a polar residue such as histidine or tyrosine (SI Fig. 10G, F, I) (10, 20). A green-absorbing state is formed when this interaction is absent in red/green CBCRs such as PPHK and Slr1393 (60) (SI Fig. 10E,F). Similarly, when an Arg residue is positioned next to the bilin NH moieties in the bilin lyase CpcT structure that also binds an *all-Z,syn* PCB (Fig. 3A), CpcT-PCB exhibits a blue color (56).

In a recent computational study that explicitly models the hydrogen bond between Asp and a *15Ea* chromophore in bacteriophytochromes, the excited-state calculations recapitulated the redshift between the Pr and Pfr states (61). 2551g3 engages the same interaction using a very different *all-Z,syn* chromphore suggesting a similar mechanism at work for FR-CBCRs. However, this interaction alone is not sufficient to explain the redshift between the Pfr states in FR-CBCRs and Cph1 because both proteins have a PCB chromophore that interacts with a counterion residue via the ring D NH moeity (Fig. 4A) [#@XB]. They only differ in their cyclic vs. extended PCB conformation. Furthermore, a series of bilin chromophores incorporated in 4718g3 consistently exhibited a redshift in the native state relative to the respective denatured state (40) (Fig. 3B). This includes phycoerythrobilin (PEB) where the D-ring is not part of the conjugated system as a result of a saturated C15 methine bridge. And the photoproducts of FR-CBCRs also exhibited redshifts (40). Taken together, these results strongly suggest that another redshift mechanism associated with ring A is at play for FR-CBCRs.

**Figure 4.**
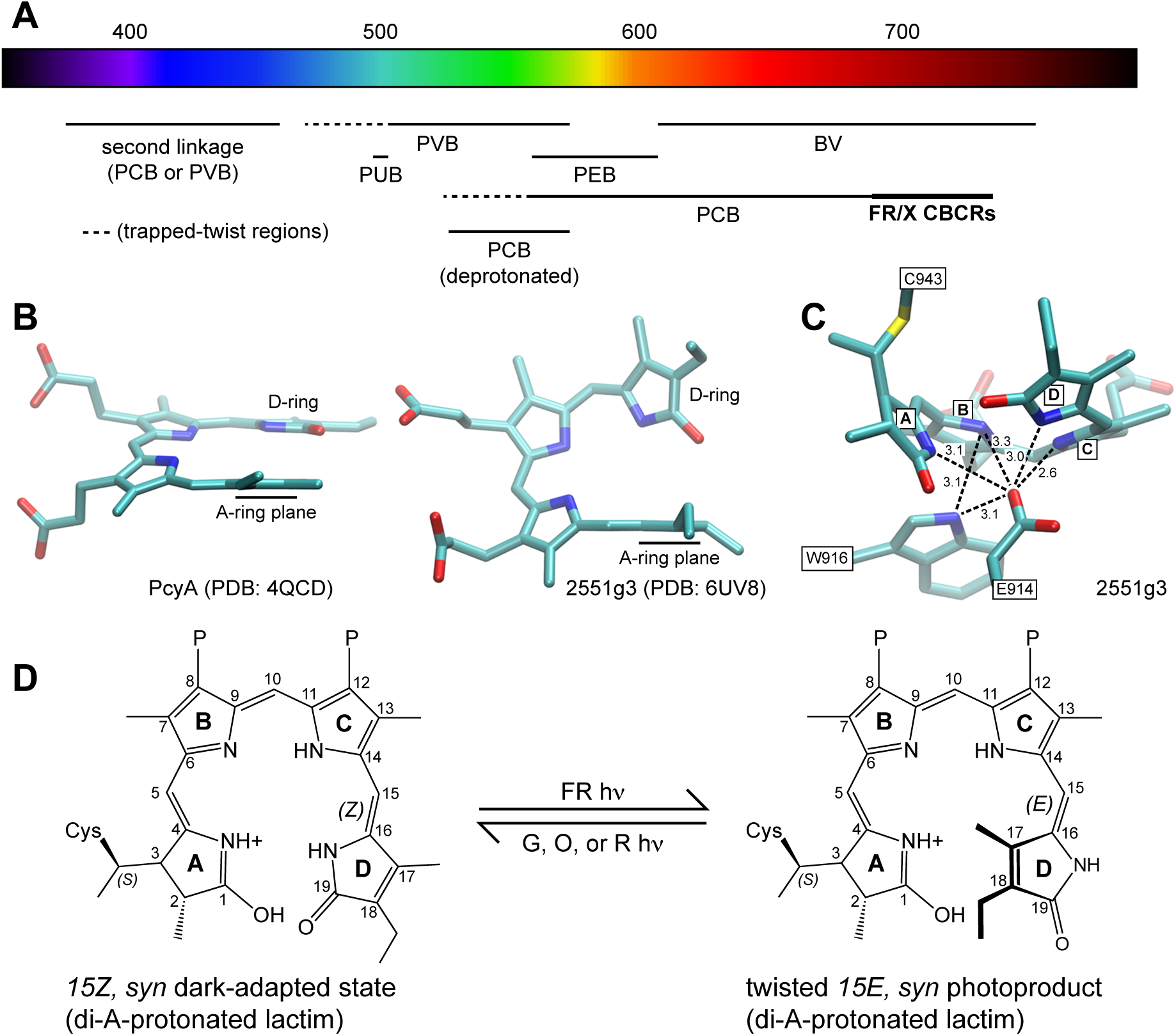
A lactim model for far-red absorption. **(A)** Chemical modifications to the bilin conjugated system resulting in a range of cyanobacterial chromophores responding to different regions of the spectrum. BV, biliverdin IXα; PCB, phycocyanobilin; PEB, phycoerythrobilin; PVB, phycoviolobilin; PUB, phycourobilin. Dashed regions indicate wavelength ranges associated with trapped-twist photoproducts in red/green and DXCF CBCRs. **(B)** The bilin A- and D-rings in PcyA (*left*) are much more coplanar than those in 2551g3 (*right*). **(C)** The PCB nitrogen atoms of 2551g3 are in a complex hydrogen bonding environmenrt, with the B-ring nitrogen interacting with the side chains of both Glu914 and Trp916. Distances are indicated in Å. **(D**) Proposed model for the 2551g3 photocycle, in which the red shift is generated by a B-deprotonated, A-di-protonated lactim tautomer that persists upon photoconversion.

Several models have been proposed to produce a redshift to the far-red region based on spectroscopic and computational studies using model compounds (62–65). Stanek and Grubmayr demonstrated that a deprotonated ring A lactim anion led to a significant red-shift in the long-wavelength band (64). Diller and colleagues proposed that far-red absorption involves a D-ring lactim tautomer (65). Krois showed that a fully protonated bilin exhibited a redshift, which was further enhanced by bringing close two carbonyls in a constrained *all-Z,syn* model compound (62). However, the Grubmayr (ring A lactim anion) model disagrees with our pH titration data on FR-CBCRs, which clearly pointed to a protonated chromophore in the Pfr state (Fig. 3). While the Diller (D-ring lactim) model supports a counterion-induced redshift, it does not explain the red-shifted spectra in photoproducts of FR-CBCRs or with PEB (Fig. 4C). The Krois (protonation) model is consistent with our pH titration results as well as the PcyA structure that have four protonated NH moieties in a more planar *all-Z,syn* BV revealed by neutron crystallography (66). While it is unclear whether a protonated, cyclic conformation of PEB would exhibit a similar redshift, this protonated state would be altered by 15,16– photoisomerization with a flipped ring D in the protein pocket. The inclusion of photocycle makes the Krois model at odds with the retention of red shift in photoproducts of FR-CBCRs (40). We also considered the A-ring lactim model because an imino ether compound mimicking the A-ring lactim produced a redshift (63). Such an A-ring lactim tautomer is not only compatible with a protonated ring system, it is also expected to revert to the lactam form upon denaturation. The A-ring lactim is expected to remain active upon 15,16-photoisomerization and still part of the conjugated system even in PEB.

We postulate that FR-CBCRs combine the Krois (protonation) effect and the Grubmayr (bilin lactim) effect to achieve a large batho-chromatic shift in an additive manner. Consistent with the Krois model, the redshift arises from protonation at the D-ring nitrogen due to interaction with the counterion residue Glu914, which evidently increases delocalization of the ring system (SI Fig. S11). In addition, a constrained cyclic bilin conformation such as the *all-Z,syn* PCB of 2551g3 would also help stabilize this kinetically mobile proton as Krois predicted (62). This effect alone may bring the PCB absorption beyond ∼700 nm, but not too much further. It is the Grubmayr effect that extends the PCB absorption to the far-red region (>720 nm). The Grubmayer effect via the A-ring lactim would also explain the red-shifts in FR-CBCRs photoproducts as well for PEB incorporated in 4718g3 (40). Although Diller et al predicted that a protonated D-ring lactim would result in a redshift of >150 nm in PCB (65), it is not known whether a protonated A-ring lactim model by itself would achieve such a large shift.

Such protonated bilin lactim have a variety of tautomers (65). To determine the most probable tautomer in the Pfr state of FR-CBCRs, we examined the *2Fo-Fc* map of 2551g3 and the protein environment of the *all-Z,syn* chromophore. We observe that the map contours connecting to Glu914 are much lower in rings B and A compared to those of rings C and D (Fig. 2A). Among four nitrogen atoms, the ring B nitrogen is also he farthest away from the counterion side chain of Glu914. It is positioned in close proximity to Trp916 on the β-face and the hydrophobic Leu944 on the α face (Fig. 2B). Given its less polar environment compared to PcyA, ring B nitrogen is most likely deprotonated in 2551g3. Rings A and D are both tilting towards the negatively charged pocket at the β-face (Fig. 2D), suggesting a di-protonated lactim tautomer at ring A or D, both of which would satisfy a protonated π system. However, the “A-ring lactim” model is deemed more favorable because it is more consistent with the spectral data that entails a redshift mechanism beyond ring D (40). This lactim tautomer is stabilized by the acidic β-facial environment in the 2551g3 structure (Fig. 2D).

We therefore propose that FR-CBCRs have a di-protonated A-ring lactim in an *all-Z,syn* chromophore with a protonated D-ring and deprotonated B-ring (Fig. 4). If the photocycles of FR-CBCRs follow the established 15,16–photoisomerization, photoconversion would yield a 15-*E,syn* PCB (Fig. 4), or less likely 15–*E,anti* conformation via a hula-twist mechanism. In either case, the A-ring lactim would be preserved after isomerization. The resulting photoproduct is expected to be red-shifted relative to the respective trapped-twist species (Fig. 3B). Compared to the “D-ring protonation” model that is shared among all canonical phytochromes, the “A-ring lactim” model represents a less unknown mechanism for spectrum tuning in the phytochrome superfamily. Collectively, these two distinct mechanisms enable PCB to absorb light in a wavelength range (∼750nm) otherwise only reachable by phytochromes utilizing BV.

### Concluding Remarks

Three major mechanisms are employed for spectrum tuning in bilin-binding proteins. *First*, proteins that use chemically distinct bilins of varying conjugated double bonds naturally absorb at different wavelengths. From PUB to BV, the longer the conjugation system, the more red-shifted the peak wavelength (SI Fig. S11A). Some CBCRs exploit a second Cys linkage to produce a blue-shifted species. *Second*, the chromophore conformation inside a protein pocket is inevitably affected by protein-chromophore interactions. Given the same bilin, a general trend of ring twist vs. peak wavelength is observed: the less ring twist confers the longer peak wavelength (Fig. 3B, SI Fig. S11B). *Third*, as we postulate in this work, far-red tuning seems to entail an additive effect from two distinct redshift mechanisms. In FR-CBCRs, “D-ring protonation” in the presence of a counterion residue works in tandem with a second mechanism via a bilin lactim tautomer. In the Pfr state of bacteriophytochromes, the biliverdin chromophore with an extra double bond is employed in addition to the characteristic protein-chromophore interaction between the highly conserved Asp residue and the D-ring NH group of the *15Ea* chromophore (Fig. 4, SI Fig. S11C). Needless to say, many aspects of the proposed far-red tuning mechanism call for future investigation by vibrational and NMR spectroscopy as well as excited state calculations.

## Materials and Methods

Detailed materials and methods are provided in the Supplementary Information. This includes information on phylogenetic analysis (67, 68), cloning, expression and purification of CBCRs, spectroscopic techniques, crystallization, and structure determination. The PDB accession codes for this work are 6UVB (100 K) and 6UV8 (room temperature).

## Acknowledgements

We thank the staff at BioCARS and LS-CAT for support in X-ray diffraction data collection. Use of the LS-CAT Sector 21 is supported by the Michigan Economic Development Corporation and the Michigan Technology Tri-Corridor under Grant 085P1000817. Use of BioCARS was also supported by the National Institute of General Medical Sciences under grant number NIH P41 GM118217. Use of the Advanced Photon Source is supported by the U. S. Department of Energy, Office of Science, Office of Basic Energy Sciences, under Contract No. DE-AC02-06CH11357. This work is supported by grants from the U. S. Department of Energy, Office of Science, Office of Basic Energy Sciences, under Contract No. E-FG02-09ER16117 to JCL, and from the National Institutes of Health under R01EY024363 to XY.

## SUPPLEMENTARY INFORMATION

### SUPPLEMENTARY METHODS

#### Bioinformatics

Candidate far-red CBCRs were identified using Anacy_4718g3 and Sta7437_1656 as queries. Phylogenetic analysis of these proteins were inferred in PhyML-Structure (1) using multiple sequence alignments generated with MAFFT (2) with the 2551g3 structure as a structural reference as previously described for FR-CBCRs and other CBCR lineages (3, 4).

#### Cloning, site-directed mutagenesis and protein purification

W911F, W916F, S917P/Q918P, and W940L variants of 2551g3 were constructed in the previously described expression construct, as were all variants of 4718g3. FR-CBCRs 01757 and 0468g3 and green/red CBCR IC523_02481 were cloned into pET28b using the procedure previously described for 2551g3 (3). For all other expression constructs, the DNA coding sequence encoding the third GAF domain of Anacy2551 from *Anabaena cylindrica* PCC 7122 (aa. 840-1026) was synthesized according to the published sequence by Genscript and subcloned into a pET24b vector using NdeI and XhoI restriction sites. A further truncated construct (denoted 2551g3), consisting of aa. 840-1019, was then sub-cloned into the pET24b vector using the same sites and was used for all other studies. All constructs carry a 6xHistidine affinity tag at the C-terminus. Site-directed mutagenesis was carried out using the standard QuikChange protocol.

4718g3 wild-type protein and mutants were expressed and purified as reported, as were W911F, W916F, S917P/Q918P, and W940L variants of 2551g3 and CBCRs 01757, 0468g3, and IC523_02481 [#@44]. All other 2551g3 constructs and mutants were co-expressed in the *E. coli* BL21(pLys) strain with a plasmid carrying heme oxygenase HO1 and bilin reductase PcyA genes to produce phycocyanobilin (PCB) (5). When the cell culture reached OD600 ∼0.4-0.6, protein over-expression was induced by addition of 1mM Isopropyl β-D-1-thiogalactopyranoside (IPTG) and 0.5 mM delta-aminolevulinic acid (ALA) followed by overnight incubation at 18°C. Cells were harvested by centrifugation, and cell pellet was kept at –80°C until purification. After cell lysis in buffer containing 20mM Tris, 50mM NaCl (pH 8.0) by sonication, the supernatant was clarified by Talon Co^2+^ affinity chromatography followed by anion exchange chromatography (5-mL HiTrap HP Q column, GE Healthcare). The fractions with good chromophore incorporation were further purified using size exclusion chromatography (10/300 GL Superdex 200 Increase, GE Healthcare). Purified protein was concentrated to 3 mg/mL. Flash frozen protein aliquots were stored in -80°C until use.

#### Crystallization and structure determination at 100 K

2551g3 in the Pfr state was crystallized following pre-illumination with filtered green light (550±20 nm; Newport) for 15 minutes before setting up crystallization. The Pfr crystals grew in the dark at room temperature using the hanging drop vapor diffusion method, in which the protein solution (3 mg/ml) and crystallization solution (17% PEG 10000, 0.1 M ammonium acetate, 0.1 M Bis-Tris pH 5.5 with Yttrium (III) chloride as an additive) were mixed in a 1:1 ratio. Green Pfr crystals of a typical size 50×50×100 (µm) appeared in 2-3 days and were harvested and cryo-protected under green safety light (500±20 nm).

The monochromatic X-ray diffraction datasets at 100 K were collected at the LS-CAT beam stations of the Advanced Photon Source, Argonne National Laboratory. All diffraction images were indexed, integrated and scaled using HKL2000 (6). Initial phases were obtained for the Pfr structure in the P4_2_22 space group by molecular replacement (PHASER) using a homologous CBCR structure (PDB: 4GLQ) as a search model (7). The final model of the Pfr structure at 100 K was refined to 2.95 Å resolution with a final R-factor and free R-factor of 0.258 and 0.303, respectively (PHENIX) (8). Coot (9) was used for all model building; illustrations were prepared using VMD and Pymol (10, 11).

#### Room-temperature serial crystallography

The 2551g3 crystals are highly susceptible to light exposure and X-ray radiation damage. Cryo-freezing also induced lattice distress likely due to the high solvent content (>70%). These factors severely limited the diffraction resolution and map quality in cryo-crystallography data. To address these problems, we carried out *in situ* X-ray diffraction experiments at room temperature using a room temperature serial crystallography platform developed by our laboratory [#@54]. From >800 crystals grown and diffracted in situ on eight crystal-on-crystal devices, we obtained a room temperature Laue diffraction dataset at the BioCARS 14-ID-B beamline of the Advanced Photon Source. Using the cryo-structure of 2551g3 as an initial model, this room temperature dataset allowed us to determine the crystal structure of 2551g3 in the Pfr state at 2.7 Å resolution. All Laue diffraction images were indexed, refined, integrated and scaled using the software package Precognition/EpinormTM (Renz Research Inc), and the final structure was refined to the final R-factor and R-free of 0.19 and 0.25, respectively (Phenix) [#69]. In our serial crystallography protocol, one crystal or one fresh crystal segment only contributed one “damage-free” diffraction image. A Laue dataset was completed by combining images from a large number of randomly orientated crystals. All details regarding this new technology platform including the automated crystal recognition, beamline control, data collection and Laue data processing protocols are reported in a separate paper.

#### UV/visible spectroscopy in solution and single crystals

Absorption spectra of 01757, 0468g3, 4718g3 variants, and W911F, W916F, S917P/Q918P, and W940L variants of 2551g3 were recorded on a Cary 50 spectrophotometer at 25°C. Photoconversion of these proteins was triggered by 728 nm LEDs (Sanyo) or a 75W xenon lamp filtered using 550 nm, 580 nm, or 600 nm band-pass filters (CVI, Chroma) (3). Solution UV/visible absorption spectra of other 2551g3 constructs were measured using Shimadzu UV-2600 spectrophotometer at room temperature. In a pH titration experiment, samples were prepared in the Pfr state under a filtered light (550±20 nm) by diluting concentrated purified proteins in a series of buffer solutions at different pHs prior to spectral measurements. Time series of absorption spectra in solution and in single crystals were recorded using a micro-spectrophotometer under pump light under band-pass filters of 550±20 or 750±20 nm (Newport). This micro-spectrophotometer is equipped with an optical lens system with 100X magnification coupled to a high-sensitivity spectrometer (QEPro, Ocean Optics), which enables accurate measurements from a sample (solution or single-crystal) with an optical surface as small as 25 µm.

## SUPPLEMENTAL FIGURE LEGENDS

**Figure S1.**
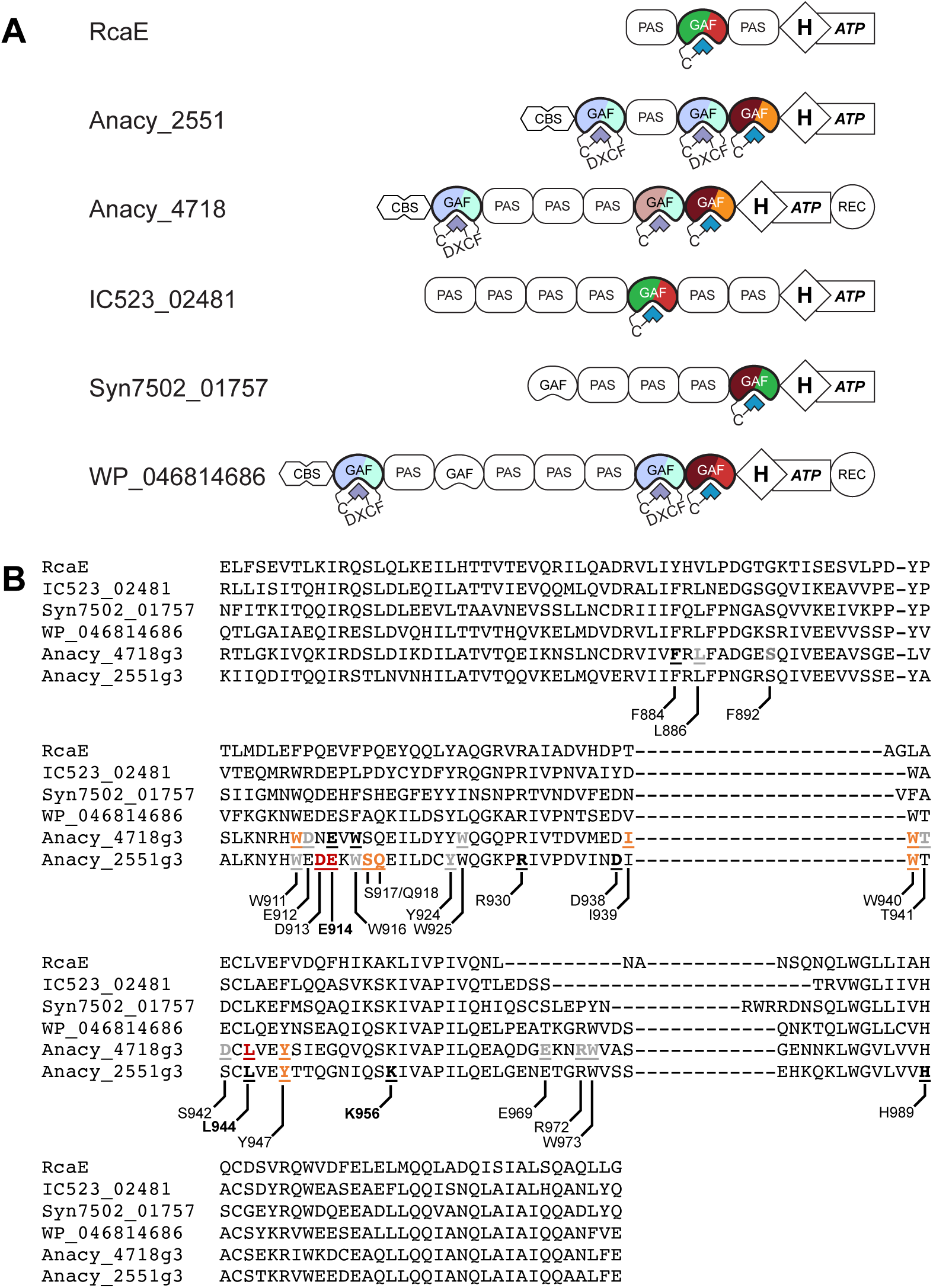
CBCRs used in this study. **(A)** Jellybean domain diagrams for proteins in this study, with the green/red CBCR RcaE shown for comparison. Characterized CBCR domains are shown in full color; predicted but uncharacterized CBCRs are shown in pastel and belong to either the DXCF or the red/green lineages [#@24,25]. Domain abbreviations: CBS, cystathionine b-synthase; GAF, cGMP phosphodiesterases, adenylyl cyclases and FhlA; H-ATP, histidine kinase bidomain; PAS, Per/Arnt/Sim; REC, response regulator receiver domain. **(B)** Characterized CBCR domains from panel A were excised from the larger sequence alignment used for phylogenetic analysis (SI Fig. S8). Amino acids targeted for site-directed mutagenesis in Anacy_2551g3 or Anacy_4718g3 are **<u>bold underlined</u>** and color-coded by phenotype using the scheme of SI Fig. S7, with numbering for 2551g3 indicated beneath. Bold numbering indicates residues conserved as the protochromic triad of RcaE [#@2].

**Figure S2.**
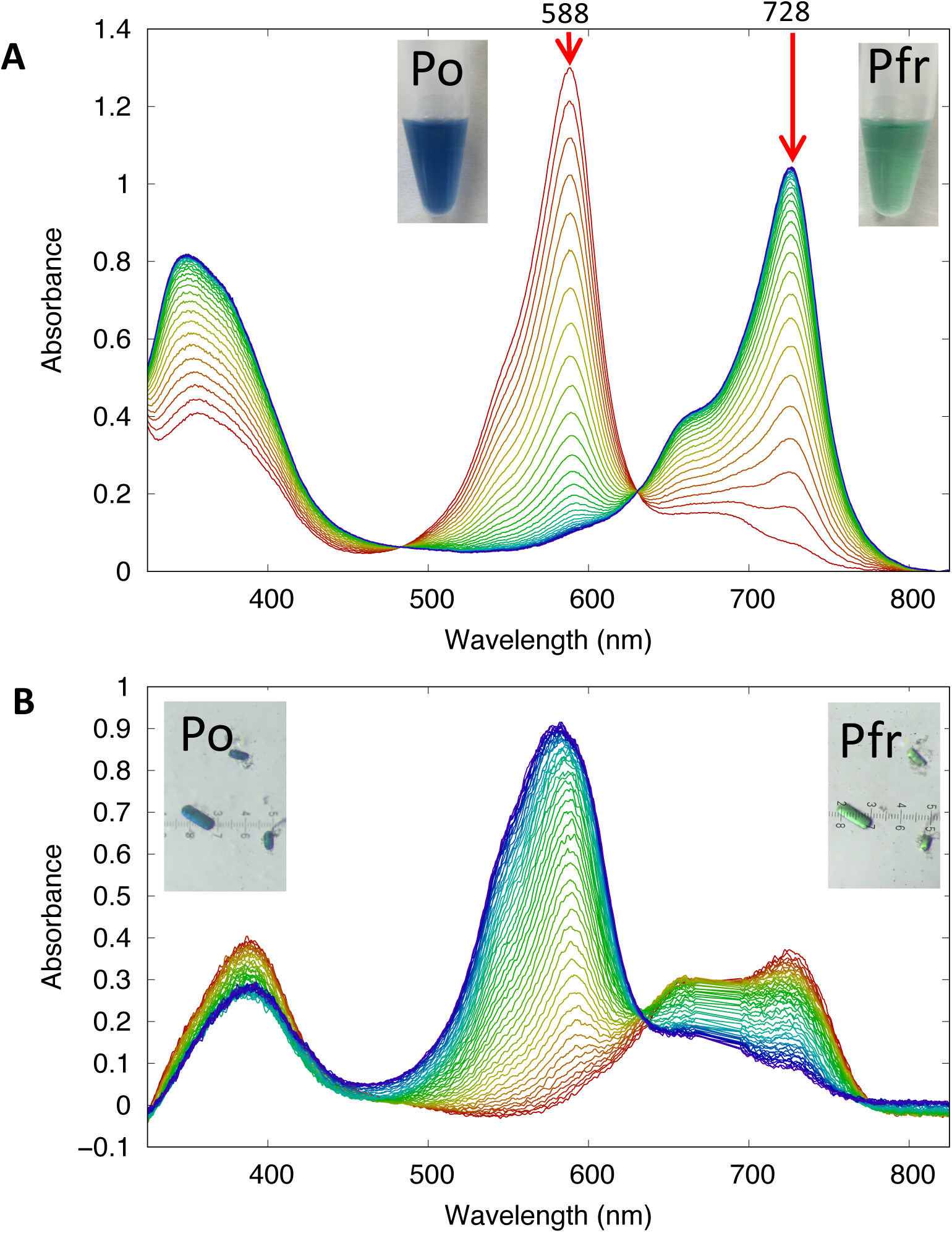
Reversible Pfr/Po photoconversion of 2551g3 in solution and single crystal. (**A)** The time course of Po→Pfr photoconversion (from red to blue) of 2551g3 in solution during a 15-second continuous illumination under filtered light (550±20 nm). (**B)** The time course of the Pfr→Po photoconversion (from red to blue) in a single crystal of 2551g3 under continuous filtered light (750±20 nm).

**Figure S3.**
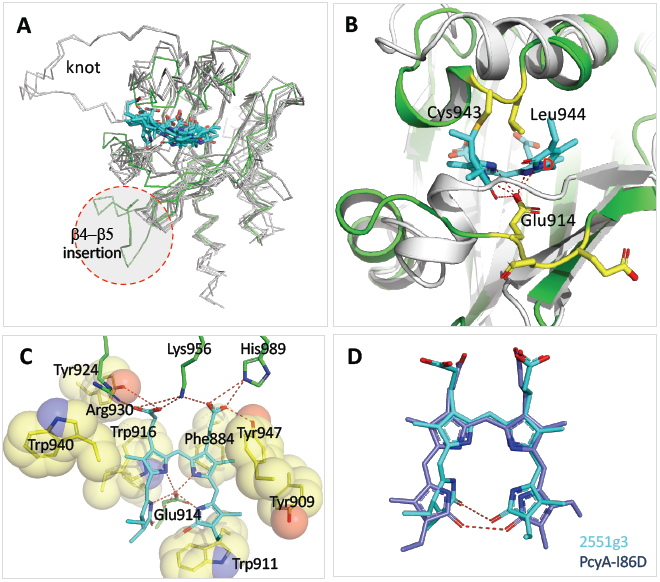
Structural comparison between 2551g3 and representative phytochromes. **(A)** Alignment of 2551g1 and representative GAF structures shows their overall agreement in protein backbone. The bilin chromophores (cyan sticks) assume similar positions in the protein pocket. 2551g3 (green wire) is unique in featuring a large insertion (b4-b5 insertion). However, 2551g3 lacks the knot structure typical for canonical phytochromes. (**B)** At the a-face, hydrophobic Leu944 points towards PCB, resulting in tilting of both rings A and D towards the b-face. The acidic side chain of Glu914 forms hydrogen bonds (red dashed lines) with all four pyrrole nitrogen atoms. AnPixJ (3W2Z; grey) is shown as a reference to highlight the differences. **C)** The *all-Z,syn* PCB in 2551g3 is stabilized by surrounding bulky residues (yellow spheres) and hydrogen-bonds (red dashed line). **D**) Superposition of the bilin chromophores in 2551g3 (cyan) and PcyA-I86D (5B4H; green) shows the close apposition between the carbonyls of rings A and D.

**Figure S4.**
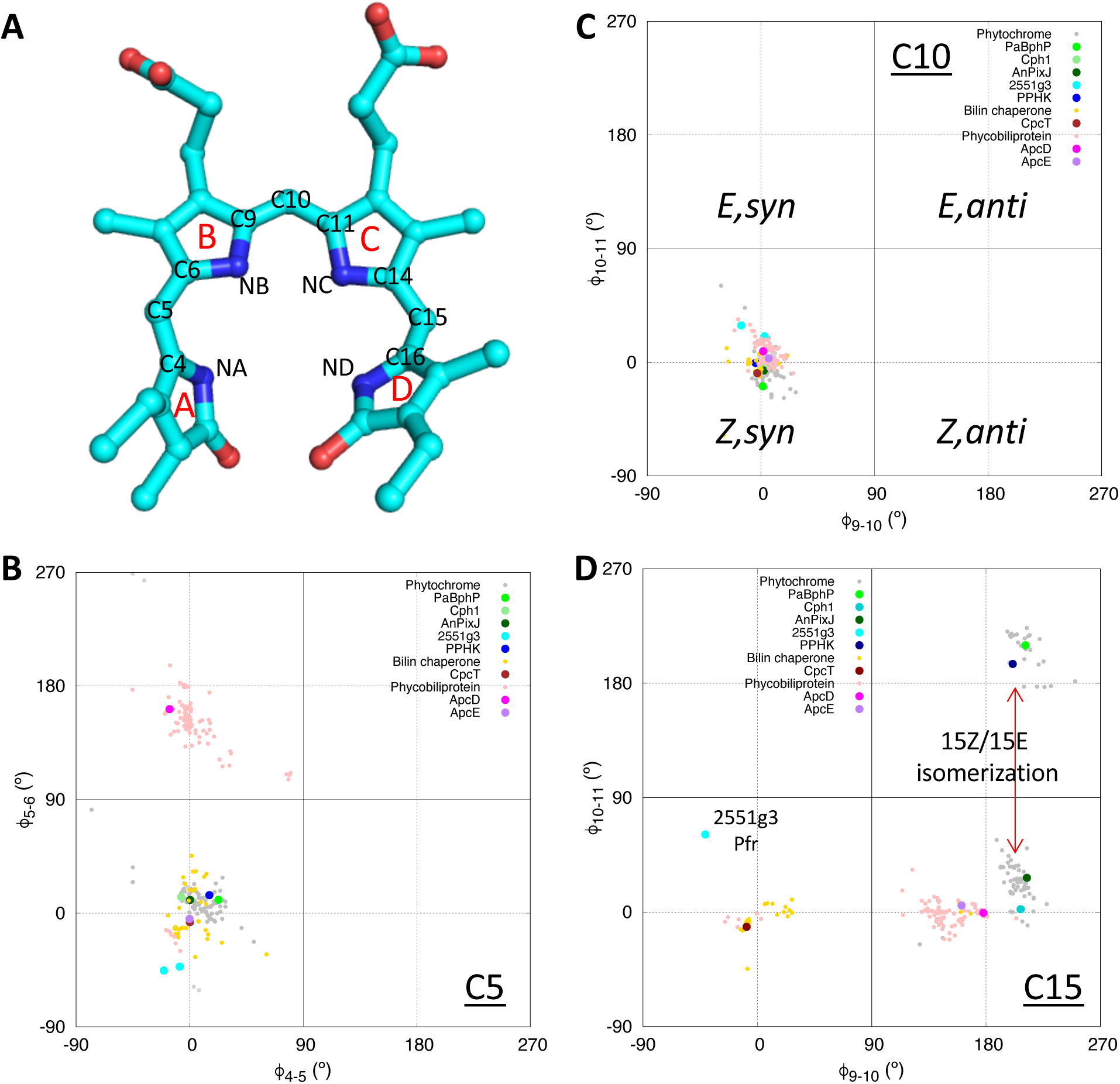
The bilin conformation distribution among bilin-binding proteins. This PDB survey examines the bilin conformations observed in 156 bilin-binding structures including the phytochrome superfamily, phycobiliproteins and bilin lyases. A total of 241 bilin chromophores (67 CYC, 61 BLA and 28 LBV; covalent or non-covalent) are used. **A)** Phycocyanobilin (PCB) of the Pfr structure of 2551g3 is shown to illustrate the atom naming. **B-D)** Scatter plots of dihedral angles in the C5, C10 and C15 methine bridges, inspired by the Ramachandran plot for depicting protein backbone structures. The bilin conformation (***Z*** vs. ***E*** and ***syn*** vs. ***anti***) in each methine bridge corresponds to a dot in one of four quadrants as defined in panel B. The dihedral angles are defined as followed. C5 methine bridge (**B**) f(4-5): N_A_-C4-C5-C6 versus f(5-6): C4-C5-C6-N_B_; C10 methine bridge (**C**) f(9-10): N_B_-C9-C10-C11 versus f(10-11): C9-C10-C11-N_C_; C15 methine bridge (**D**) f(14-15): N_C_-C14-C15-C16; f(15-16): C14-C15-C16-N_D_.

**Figure S5.**
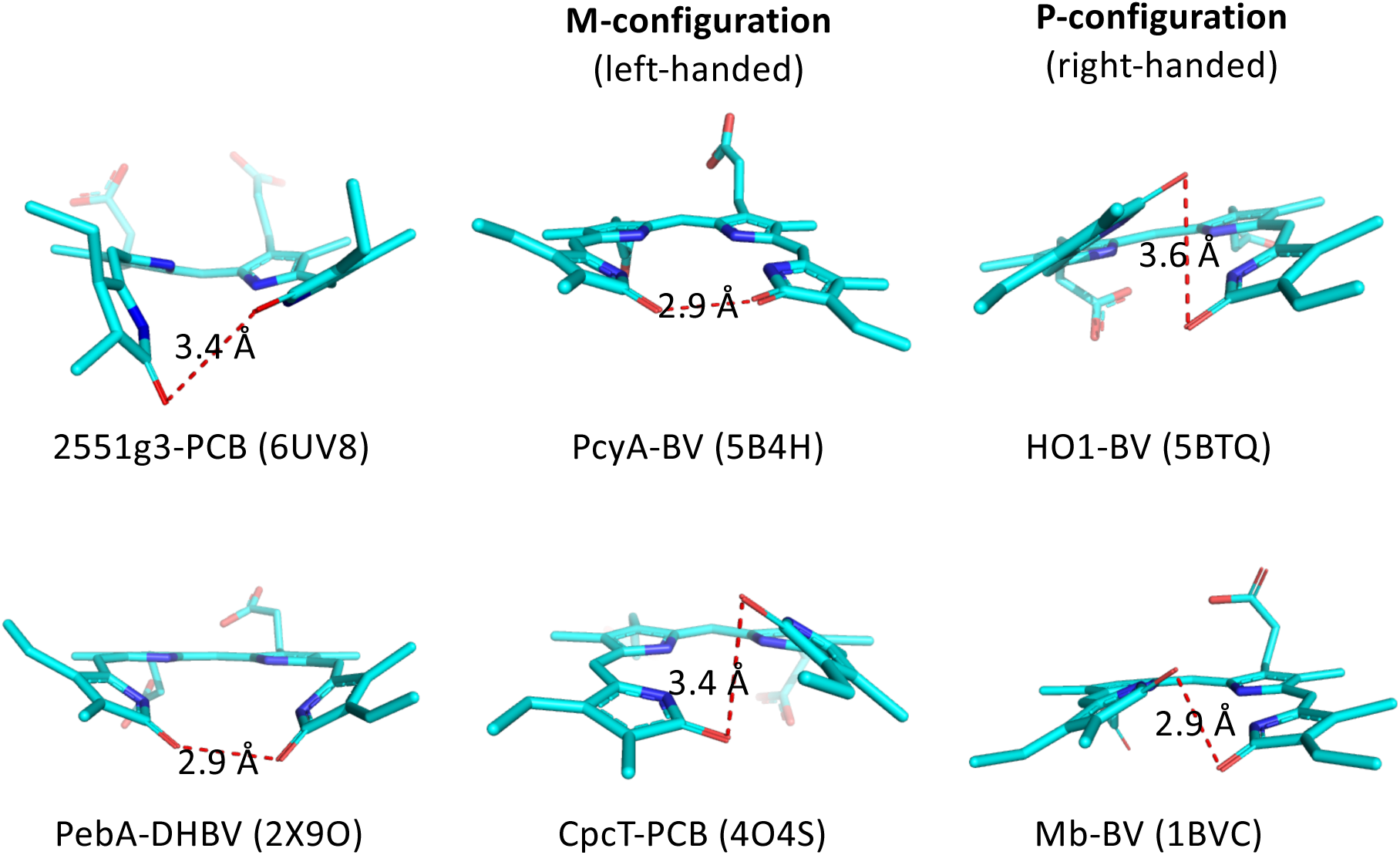
Representative *all-Z,syn* bilins in the Protein Data Bank. Among the *all-Z,syn* bilins, only PCB in 2551g3 forms a covalent linkage to the protein moiety via a cysteine anchor. The majority of *all-Z,syn* bilins adopt an axial chirality in a left-handed configuration (denoted M-configuration) as shown in biliverdin reductase PcyA and bilin lyase CpcT. In a few cases as in human heme oxygenase HO1 and myogloblin, the bilin chromophore adopts a right-handed P-configuration. Distances between the carbonyl oxygen atoms of rings A and D are shown along with the red dashed lines.

**Figure S6.**
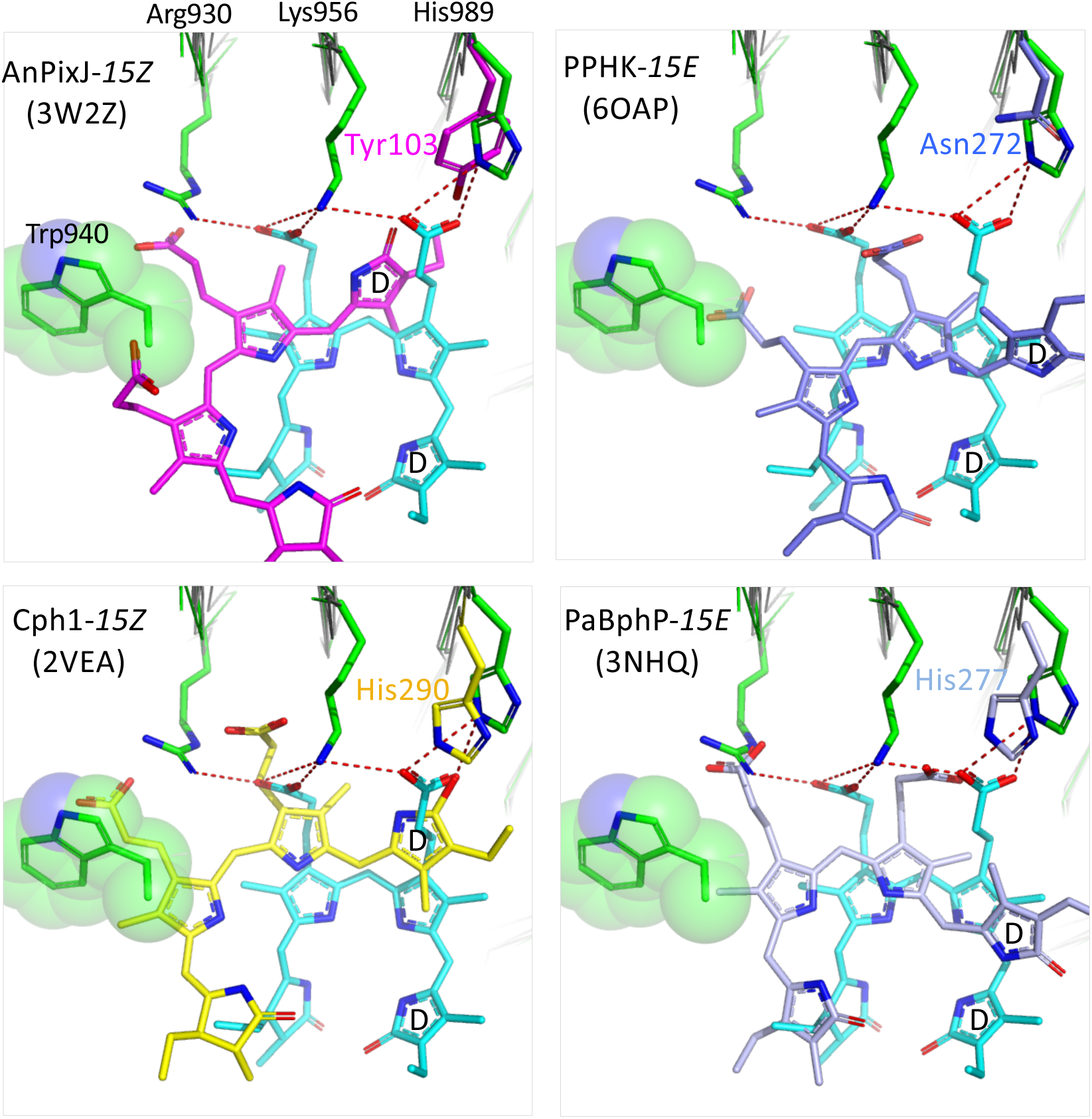
The 2551g3-Pfr structure shows an unusual chromophore disposition compared to the representative phytochrome structures. All structures are aligned according to the protein framework in the GAF core. The bilin chromophore and protein side chains of 2551g3 are colored in cyan and green, respectively. Green spheres marks the bulky side chain of Trp940 in 2551g3, which would impose a steric clash with the ring B propionate of the bilin chromophore in all other structures.

**Figure S7.**
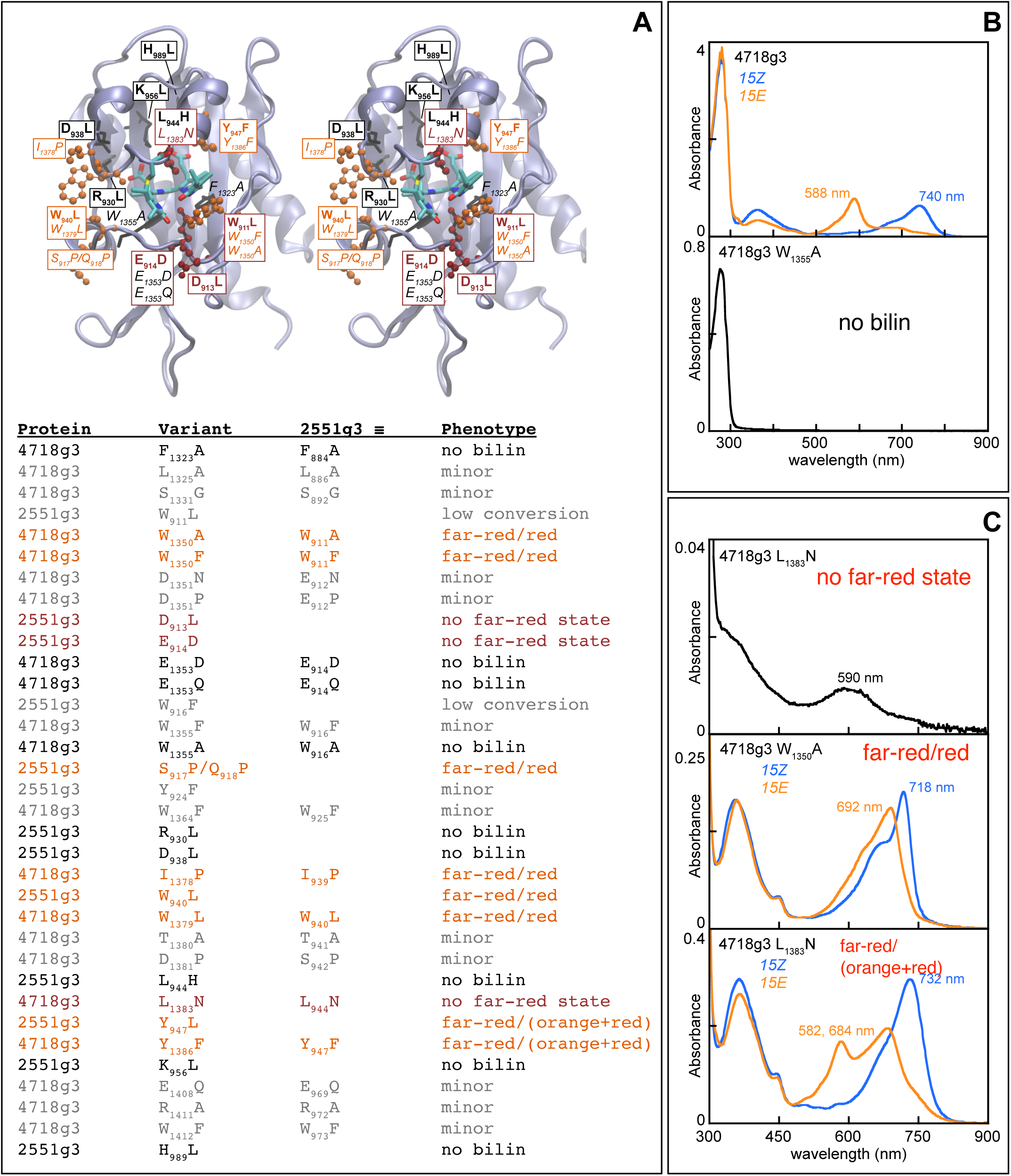
Analysis of variant proteins. **(A)** A stereo view of the 2551g3 structure is shown. amino acid substitutions giving rise to significant phenotypes are indicated on the structure (black, no bilin; orange, red-shifted photoproduct; brick red, loss of far-red absorption). All variants tested are listed beneath the structures, with those yielding only minor phenotypes in grey. **(B)** Wild-type 4718g3 (top: *15Z*, blue; *15E*, bottom) is compared to a representative example which lost bilin binding (W_1355_A 4718g3). **(C)** Representative examples are shown for loss of far-red absorption (L_1383_N 4718g3, top), red-absorbing photoproduct (W_1355_A 4718g3, middle), and mixed photoproduct species (W_1355_A 4718g3, bottom), with the *15Z* state in blue and the *15E* state in orange. L_1383_N (black) also exhibited poor chromophorylation, so chemical configuration could not be assigned.

**Figure S8.**
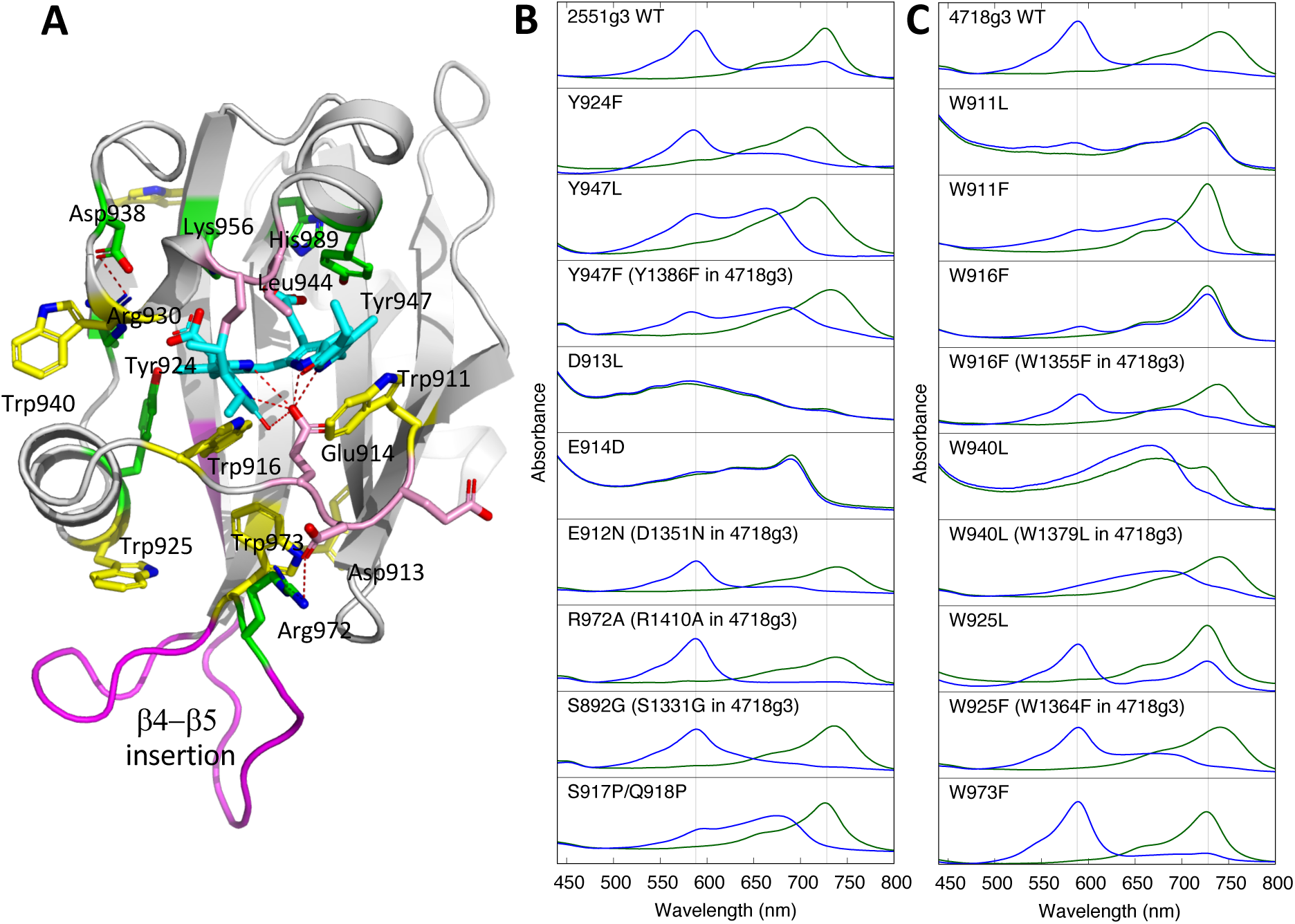
Mutational studies in FR-CBCRs. A) Sites of mutagenesis. The room temperature Pfr structure (PDBID: 6UV8) is used for illustration. The PCB chromophore is colored in cyan. Residues in green are those that directly interact with the propionates. Residues in pink engage close contacts to the a- and b-faces of PCB. The Trp residues are colored in yellow. B) Absorption spectra of WT 2551g3 and mutants of 2551g3 and 4718g3. C) Absorption spectra of WT 4718g3 and single mutants of selected Trp residues. Labels in parentheses indicate that the spectra were obtained from 4718g3 along with the equivalent mutations in 2551g3. Green and blue curves represent the *15Z* and *15E* states following 15-minute illumination of filtered 550 and 750 nm, respectively. The peak positions of the Pfr (728 nm) and Po (588 nm) states of 2551g3 are marked by grey thin lines.

**Figure S9.**
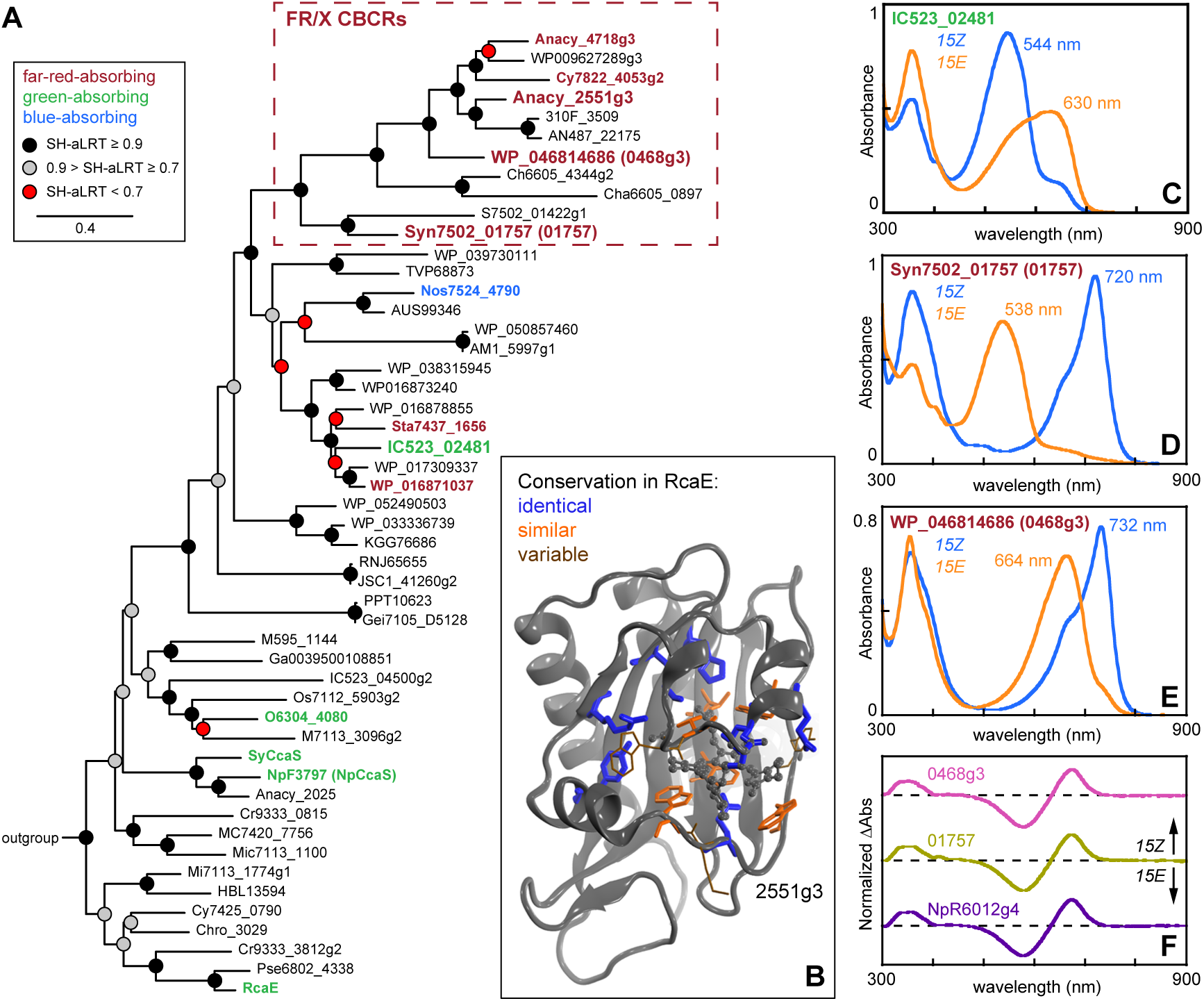
Identification and characterization of additional FR-CBCRs. **(A)** A maximum-likelihood phylogenetic tree is shown for CBCRs in the GGR lineage, with the knotless phytochrome outgroup omitted for clarity. Characterized proteins are in **bold** and are color-coded by *15Z* absorption maximum. The FR/X cluster identified by this analysis is indicated. **(B)** The 2551g3 structure is shown, with residues proximal to the chromophore color-coded by conservation in the green/red CBCR RcaE as indicated. **(C)** Absorption spectra are shown for IC523_02481 in the *15Z* (blue) and *15E* (orange) states. **(D)** Absorption spectra are shown for 01757 in the color scheme of panel C. **(E)** Absorption spectra are shown for 0468g3 in the color scheme of panel C.

**Figure S10.**
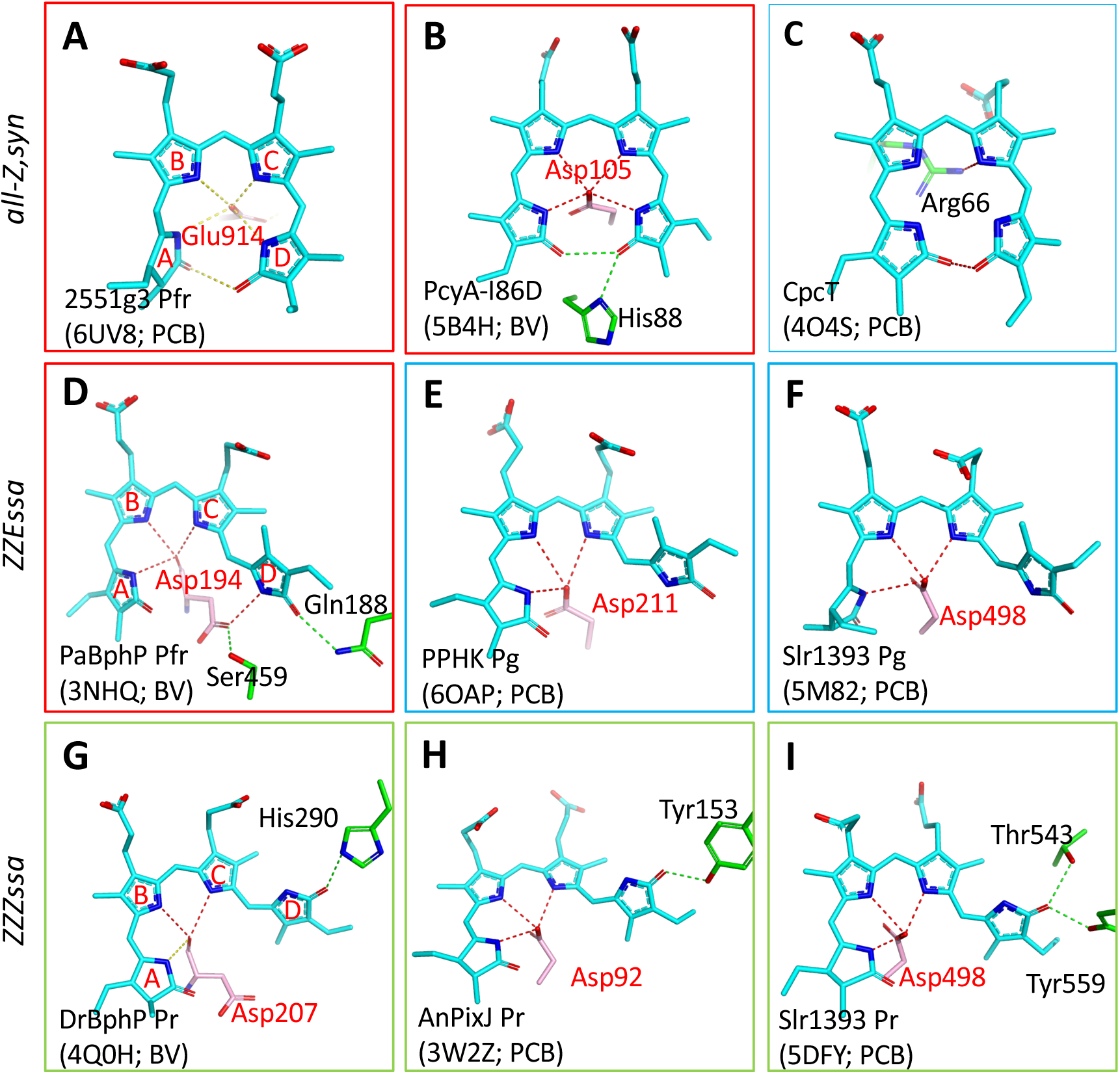
Protein-chromophore interactions in bilin-binding proteins. Direct interactions between the protein moiety and pyrrole rings within a distance of 3.5 Å are shown in dashed lines for nine representative bilin-binding structures organized according to their bilin conformations. Each panel is annotated with the bilin type (PCB or BV; in cyan) along with the corresponding PDB code. In all FR-conferring structures (A, B, D; in red outline), the ring D nitrogen directly interacts with the acidic side chain of Glu or Asp (pink) while the ring D carbonyl is hydrogen bonded to a polar moiety (green dashed lines). The green-absorbing phenotype correlates with the absence of the corresponding acidic residue near ring D lactam (C, E, F; in blue outline). In the red-absorbing Pr state, the ring D carbonyl is stabilized by a polar moiety while no counterion is found for the ring D nitrogen (G, H and I; in green outline).

**Figure S11.**
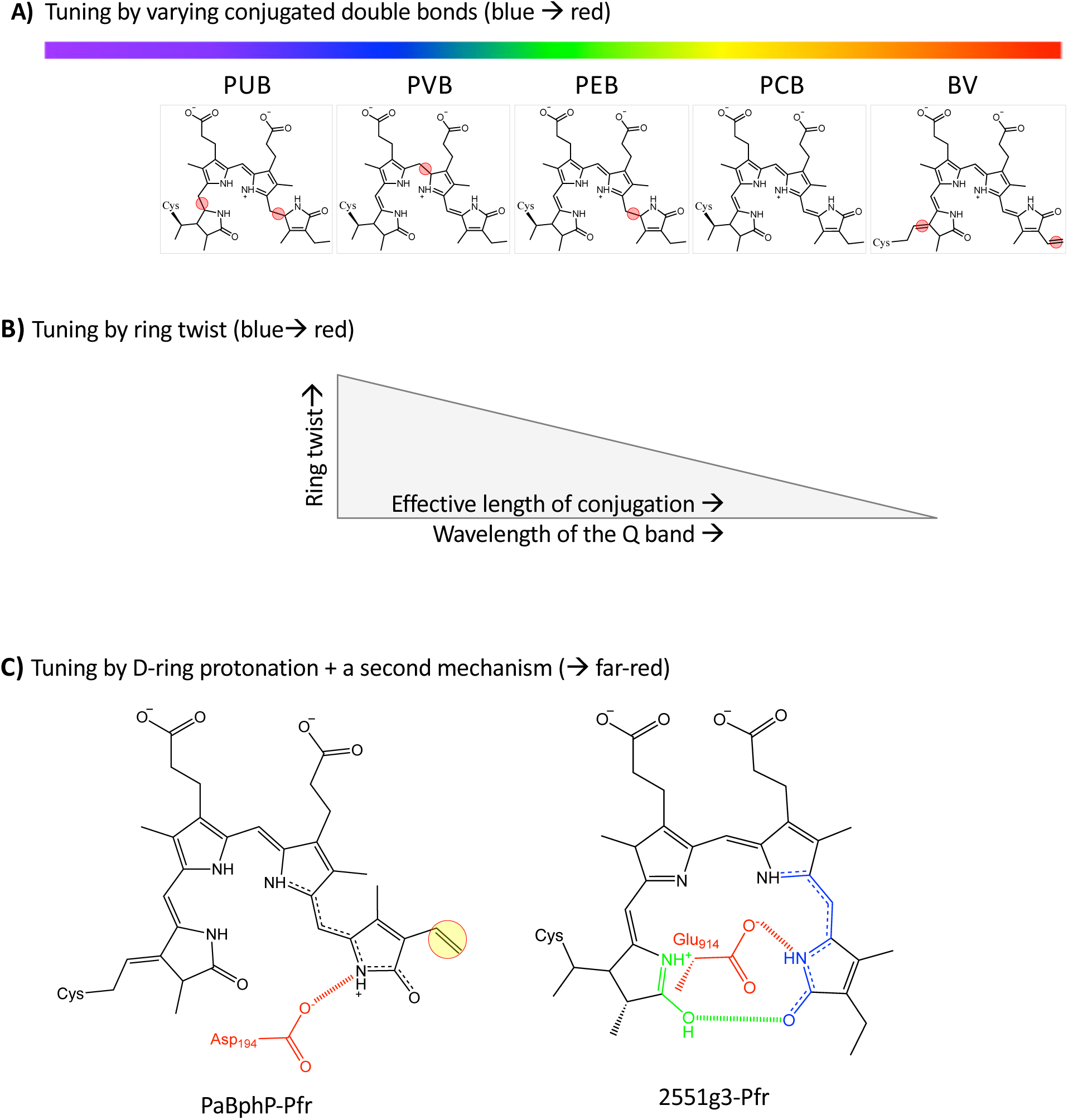
Spectrum tuning mechanisms in bilin-binding proteins. **A)** Proteins that use chemically distinct bilins of varying conjugated double bonds naturally absorb at different wavelengths. From PUB to BV, the longer the conjugation system, the more red-shifted the peak wavelength. Red circles mark those bonds that differ from the reference phycocyanobilin (PCB). **B)** The chromophore conformation inside a protein pocket is affected by protein-chromophore interactions. Given the same bilin, a general trend of ring twist vs. peak wavelength is observed: the less ring twist confers the longer peak wavelength. **C)** Far-red tuning may entail two mechanisms via an additive effect. The redshift via D-ring protonation in the presence of a counterion residue (Asp or Glu; in red) works in tandem with a second redshift mechanism either via a bilin lactim tautomer as in FR-CBCRs or an extra double bond as in bacteriophytochromes.

---

**Table S1.** Crystallography Data Collection and Refinement Statistics.

|  |  |  |
| --- | --- | --- |
| Structure | 2551g3-RT (Pfr) | 2551g3-cryo (Pfr) |
| Space group | P4<sub>2</sub>22 | P4<sub>2</sub>22 |
| Cell parameters |  |  |
| *a, b, c* (Å) | 108.8, 108.8, 67.9 | 108.8, 108.8, 68.7 |
| $\alpha, \beta, \gamma$ (°) | 90, 90, 90 | 90, 90, 90 |
| Residues | 839-1018 | 839-1018 |
| Chain | A | A |
| Ligands | 1 CYC / 7 waters | 1 CYC / 0 waters |
| Diffraction data |  |  |
| X-ray source | 14-ID-B, APS | 21-ID-D, APS |
| Methods | Laue<br>Serial | Single-wavelength<br>1° oscillation |
|  | Room temperature | 100 K |
| Number of crystals | 800<br>50-2.7 (3.31-2.70) | 1<br>50-2.90 (2.95-2.90) |
| Resolution (Å) | 0.038 (0.746) | 0.033 (0.705) |
| *R*<sub>merge</sub> | 98.7 (99.8) | 98.9 (100.0) |
| Completeness (%) | 10.3 (8.5) | 6.7 (6.5) |
| Redundancy | 43.4 (1.3) | 30.2 (1.0) |
| *I*/$\sigma$(*I*) | | |
| Refinement |  |  |
| Resolution (Å) | 47.73-2.70 (2.79-2.70) | 39.3-2.95 (3.01-2.95) |
| *R* | 0.180 (0.312) | 0.242 (0.361) |
| *R*<sub>free</sub> | 0.242 (0.293) | 0.298 (0.303) |
| RMSD |  |  |
| Bond length (Å) | 0.008 | 0.008 |
| Bond angle (°) | 1.231 | 1.248 |
| Average *B* (Å<sup>2</sup>) | 66.58 | 98.9 |
| Ramachandran |  |  |
| Favored (%) | 91.76 | 96.1 |
| Allowed (%) | 8.24 | 3.3 |
| Disallowed (%) | 0.00 | 0.55 |
| PDB entry | 6UV8 | 6UVB |

## Notes

### Competing Interest Statement

The authors have declared no competing interest.

## References

1. M. Ikeuchi, T. Ishizuka, Cyanobacteriochromes: a new superfamily of tetrapyrrole-binding photoreceptors in cyanobacteria. Photochem. Photobiol. Sci. 7, 1159 (2008).

2. Y. Hirose, et al., Green/red cyanobacteriochromes regulate complementary chromatic acclimation via a protochromic photocycle. Proc. Natl. Acad. Sci. 110, 4974–4979 (2013).

3. F. Gan, et al., Extensive remodeling of a cyanobacterial photosynthetic apparatus in far-red light. Science 345, 1312–1317 (2014).

4. K. Fushimi, R. Narikawa, Cyanobacteriochromes: photoreceptors covering the entire UV-to-visible spectrum. Curr. Opin. Struct. Biol. 57, 39–46 (2019).

5. N. C. Rockwell, Y.-S. Su, J. C. Lagarias, Phytochrome Structure and Signaling Mechanisms. Annu. Rev. Plant Biol. 57, 837–858 (2006).

6. C. Song, et al., Two ground state isoforms and a chromophore D-ring photoflip triggering extensive intramolecular changes in a canonical phytochrome. Proc. Natl. Acad. Sci. 108, 3842–3847 (2011).

7. R. Narikawa, et al., Structures of cyanobacteriochromes from phototaxis regulators AnPixJ and TePixJ reveal general and specific photoconversion mechanism. Proc. Natl. Acad. Sci. 110, 918–923 (2013).

8. E. S. Burgie, J. M. Walker, G. N. Phillips Jr., R. D. Vierstra, A Photo-Labile Thioether Linkage to Phycoviolobilin Provides the Foundation for the Blue/Green Photocycles in DXCF-Cyanobacteriochromes. Structure 21, 88–97 (2013).

9. X. Yang, Z. Ren, J. Kuk, K. Moffat, Temperature-scan cryocrystallography reveals reaction intermediates in bacteriophytochrome. Nature 479, 428–432 (2011).

10. X. Xu, et al., Structural elements regulating the photochromicity in a cyanobacteriochrome. Proc. Natl. Acad. Sci. 117, 2432–2440 (2020).

11. S. Lim, et al., Correlating structural and photochemical heterogeneity in cyanobacteriochrome NpR6012g4. Proc. Natl. Acad. Sci. 115, 4387–4392 (2018).

12. K. A. Franklin, P. H. Quail, Phytochrome functions in Arabidopsis development. J. Exp. Bot. 61, 11–24 (2010).

13. J. J. Casal, Photoreceptor Signaling Networks in Plant Responses to Shade. Annu. Rev. Plant Biol. 64, 403–427 (2013).

14. X. Yang, J. Kuk, K. Moffat, Crystal structure of Pseudomonas aeruginosa bacteriophytochrome: Photoconversion and signal transduction. Proc. Natl. Acad. Sci. 105, 14715–14720 (2008).

15. G. Enomoto, Ni-Ni-Win, R. Narikawa, M. Ikeuchi, Three cyanobacteriochromes work together to form a light color-sensitive input system for c-di-GMP signaling of cell aggregation. Proc. Natl. Acad. Sci. 112, 8082–8087 (2015).

16. Y. Hirose, et al., Diverse Chromatic Acclimation Processes Regulating Phycoerythrocyanin and Rod-Shaped Phycobilisome in Cyanobacteria. Mol. Plant 12, 715–725 (2019).

17. L. B. Wiltbank, D. M. Kehoe, Diverse light responses of cyanobacteria mediated by phytochrome superfamily photoreceptors. Nat. Rev. Microbiol. 17, 37–50 (2019).

18. G. Gourinchas, et al., Long-range allosteric signaling in red light–regulated diguanylyl cyclases. Sci. Adv. 3, e1602498 (2017).

19. L.-O. Essen, J. Mailliet, J. Hughes, The structure of a complete phytochrome sensory module in the Pr ground state. Proc. Natl. Acad. Sci. 105, 14709–14714 (2008).

20. J. R. Wagner, J. S. Brunzelle, K. T. Forest, R. D. Vierstra, A light-sensing knot revealed by the structure of the chromophore-binding domain of phytochrome. Nature 438, 325–331 (2005).

21. E. S. Burgie, J. Zhang, R. D. Vierstra, Crystal Structure of Deinococcus Phytochrome in the Photoactivated State Reveals a Cascade of Structural Rearrangements during Photoconversion. Structure 24, 448–457 (2016).

22. N. C. Rockwell, S. S. Martin, K. Feoktistova, J. C. Lagarias, Diverse two-cysteine photocycles in phytochromes and cyanobacteriochromes. Proc. Natl. Acad. Sci. 108, 11854–11859 (2011).

23. N. C. Rockwell, S. S. Martin, A. G. Gulevich, J. C. Lagarias, Phycoviolobilin Formation and Spectral Tuning in the DXCF Cyanobacteriochrome Subfamily. Biochemistry 51, 1449–1463 (2012).

24. N. C. Rockwell, S. S. Martin, J. C. Lagarias, Red/Green Cyanobacteriochromes: Sensors of Color and Power. Biochemistry 51, 9667–9677 (2012).

25. R. Narikawa, G. Enomoto, Ni-Ni-Win, K. Fushimi, M. Ikeuchi, A New Type of Dual-Cys Cyanobacteriochrome GAF Domain Found in Cyanobacterium *Acaryochloris marina*, Which Has an Unusual Red/Blue Reversible Photoconversion Cycle. Biochemistry 53, 5051–5059 (2014).

26. K. Fushimi, et al., Cyanobacteriochrome Photoreceptors Lacking the Canonical Cys Residue. Biochemistry 55, 6981–6995 (2016).

27. J. J. Tabor, A. Levskaya, C. A. Voigt, Multichromatic Control of Gene Expression in Escherichia coli. J. Mol. Biol. 405, 315–324 (2011).

28. A. Hueso-Gil, Á. Nyerges, C. Pál, B. Calles, V. de Lorenzo, Multiple-Site Diversification of Regulatory Sequences Enables Interspecies Operability of Genetic Devices. ACS Synth. Biol. 9, 104–114 (2020).

29. M. Blain-Hartung, N. C. Rockwell, J. C. Lagarias, Light-Regulated Synthesis of Cyclic-di-GMP by a Bidomain Construct of the Cyanobacteriochrome Tlr0924 (SesA) without Stable Dimerization. Biochemistry 56, 6145–6154 (2017).

30. M. Blain-Hartung, et al., Cyanobacteriochrome-based photoswitchable adenylyl cyclases (cPACs) for broad spectrum light regulation of cAMP levels in cells. J. Biol. Chem. 293, 8473–8483 (2018).

31. K. Fushimi, G. Enomoto, M. Ikeuchi, R. Narikawa, Distinctive Properties of Dark Reversion Kinetics between Two Red/Green-Type Cyanobacteriochromes and their Application in the Photoregulation of cAMP Synthesis. Photochem. Photobiol. 93, 681–691 (2017).

32. P. Ramakrishnan, J. J. Tabor, Repurposing *Synechocystis* PCC6803 UirS–UirR as a UV-Violet/Green Photoreversible Transcriptional Regulatory Tool in *E. coli*. ACS Synth. Biol. 5, 733–740 (2016).

33. S. Lim, et al., Photoconversion changes bilin chromophore conjugation and protein secondary structure in the violet/orange cyanobacteriochrome NpF2163g3. Photochem. Photobiol. Sci. (2014) https://doi.org/10.1039/C3PP50442E (May 9, 2014).

34. N. C. Rockwell, S. S. Martin, J. C. Lagarias, Identification of DXCF cyanobacteriochrome lineages with predictable photocycles. Photochem. Photobiol. Sci. 14, 929–941 (2015).

35. N. C. Rockwell, S. S. Martin, J. C. Lagarias, There and Back Again: Loss and Reacquisition of Two-Cys Photocycles in Cyanobacteriochromes. Photochem. Photobiol. 93, 741–754 (2017).

36. T. Ishizuka, et al., The Cyanobacteriochrome, TePixJ, Isomerizes Its Own Chromophore by Converting Phycocyanobilin to Phycoviolobilin. Biochemistry 50, 953–961 (2011).

37. N. C. Rockwell, S. S. Martin, J. C. Lagarias, Mechanistic Insight into the Photosensory Versatility of DXCF Cyanobacteriochromes. Biochemistry 51, 3576–3585 (2012).

38. S. Osoegawa, et al., Identification of the Deprotonated Pyrrole Nitrogen of the Bilin-Based Photoreceptor by Raman Spectroscopy with an Advanced Computational Analysis. J. Phys. Chem. B 123, 3242–3247 (2019).

39. C. Zhao, F. Gan, G. Shen, D. A. Bryant, RfpA, RfpB, and RfpC are the Master Control Elements of Far-Red Light Photoacclimation (FaRLiP). Front. Microbiol. 6 (2015).

40. N. C. Rockwell, S. S. Martin, J. C. Lagarias, Identification of Cyanobacteriochromes Detecting Far-Red Light. Biochemistry 55, 3907–3919 (2016).

41. K. Fushimi, M. Ikeuchi, R. Narikawa, The Expanded Red/Green Cyanobacteriochrome Lineage: An Evolutionary Hot Spot. Photochem. Photobiol. 93, 903–906 (2017).

42. H. Scheer, X. Yang, K.-H. Zhao, Biliproteins and their Applications in Bioimaging. Procedia Chem. 14, 176–185 (2015).

43. K. D. Piatkevich, F. V. Subach, V. V. Verkhusha, Engineering of bacterial phytochromes for near-infrared imaging, sensing, and light-control in mammals. Chem. Soc. Rev. (2013) https://doi.org/10.1039/C3CS35458J (February 8, 2013).

44. J. Lecoq, M. J. Schnitzer, An infrared fluorescent protein for deeper imaging. Nat. Biotechnol. 29, 715–716 (2011).

45. G. S. Filonov, et al., Bright and stable near-infrared fluorescent protein for in vivo imaging. Nat. Biotechnol. 29, 757–761 (2011).

46. D. M. Chudakov, M. V. Matz, S. Lukyanov, K. A. Lukyanov, Fluorescent Proteins and Their Applications in Imaging Living Cells and Tissues. Physiol. Rev. 90, 1103–1163 (2010).

47. X. Shu, et al., Mammalian Expression of Infrared Fluorescent Proteins Engineered from a Bacterial Phytochrome. Science 324, 804–807 (2009).

48. O. S. Oliinyk, A. A. Shemetov, S. Pletnev, D. M. Shcherbakova, V. V. Verkhusha, Smallest near-infrared fluorescent protein evolved from cyanobacteriochrome as versatile tag for spectral multiplexing. Nat. Commun. 10 (2019).

49. K. Fushimi, et al., Photoconversion and Fluorescence Properties of a Red/Green-Type Cyanobacteriochrome AM1_C0023g2 That Binds Not Only Phycocyanobilin But Also Biliverdin. Front. Microbiol. 7 (2016).

50. K. Fushimi, et al., Rational conversion of chromophore selectivity of cyanobacteriochromes to accept mammalian intrinsic biliverdin. Proc. Natl. Acad. Sci. 116, 8301–8309 (2019).

51. G. A. Gambetta, J. C. Lagarias, Genetic engineering of phytochrome biosynthesis in bacteria. Proc. Natl. Acad. Sci. 98, 10566–10571 (2001).

52. J. Zhang, et al., Fused-Gene Approach to Photoswitchable and Fluorescent Biliproteins. Angew. Chem. Int. Ed. 49, 5456–5458 (2010).

53. Z. Ren, et al., Crystal-on-crystal chips for *in situ* serial diffraction at room temperature. Lab. Chip 18, 2246–2256 (2018).

54. N. C. Rockwell, L. Shang, S. S. Martin, J. C. Lagarias, Distinct classes of red/far-red photochemistry within the phytochrome superfamily. Proc. Natl. Acad. Sci. 106, 6123–6127 (2009).

55. Y. Hagiwara, M. Sugishima, Y. Takahashi, K. Fukuyama, Crystal structure of phycocyanobilin:ferredoxin oxidoreductase in complex with biliverdin IX, a key enzyme in the biosynthesis of phycocyanobilin. Proc. Natl. Acad. Sci. 103, 27–32 (2006).

56. W. Zhou, et al., Structure and Mechanism of the Phycobiliprotein Lyase CpcT. J. Biol. Chem. 289, 26677–26689 (2014).

57. A. Marx, N. Adir, Allophycocyanin and phycocyanin crystal structures reveal facets of phycobilisome assembly. Biochim. Biophys. Acta BBA - Bioenerg. 1827, 311–318 (2013).

58. J. Zhang, et al., Structure of phycobilisome from the red alga Griffithsia pacifica. Nature 551, 57–63 (2017).

59. Y. Hagiwara, et al., Atomic-resolution structure of the phycocyanobilin:ferredoxin oxidoreductase I86D mutant in complex with fully protonated biliverdin. FEBS Lett. 590, 3425–3434 (2016).

60. H. Shin, Z. Ren, X. Zeng, S. Bandara, X. Yang, Structural basis of molecular logic OR in a dual-sensor histidine kinase. Proc. Natl. Acad. Sci. 116, 19973–19982 (2019).

61. E. Maximowitsch, T. Domratcheva, “A Hydrogen Bond Between Linear Tetrapyrrole and Conserved Aspartate Causes the Far-Red Shifted Absorption of Phytochrome Photoreceptors” (2020) https://doi.org/10.26434/chemrxiv.12278780.v1 (June 1, 2020).

62. D. Krois, Geometry versus basicity of bilatrienes: Stretched and helical protonated biliverdins. Monatshefte Für Chem. Chem. Mon. 122, 495–506 (1991).

63. R. Micura, K. Grubmayr, LONG-WAVELENGTH ABSORBING DERIVATIVES OF PHYCOCYANOBILIN: NEW STRUCTURAL ASPECTS ON PHYTOCHROME. 6.

64. M. Stanek, K. Grubmayr, Deprotonated 2,3-Dihydrobilindiones-Models for the Chromophore of the Far-Red-Absorbing Form of Phytochrome. Chem. Eur. J 4, 1660–1666 (1998).

65. P. Singer, S. Fey, A. H. Göller, G. Hermann, R. Diller, Femtosecond Dynamics in the Lactim Tautomer of Phycocyanobilin: A Long-Wavelength Absorbing Model Compound for the Phytochrome Chromophore. ChemPhysChem 15, 3824–3831 (2014).

66. M. Unno, et al., Insights into the Proton Transfer Mechanism of a Bilin Reductase PcyA Following Neutron Crystallography. J. Am. Chem. Soc. 137, 5452–5460 (2015).

67. S. Q. Le, O. Gascuel, Accounting for Solvent Accessibility and Secondary Structure in Protein Phylogenetics Is Clearly Beneficial. Syst. Biol. 59, 277–287 (2010).

68. K. Katoh, D. M. Standley, MAFFT Multiple Sequence Alignment Software Version 7: Improvements in Performance and Usability. Mol. Biol. Evol. 30, 772–780 (2013).

## SI References

1. C. Gasser, et al., Engineering of a red-light–activated human cAMP/cGMP-specific phosphodiesterase. Proc. Natl. Acad. Sci. 111, 8803–8808 (2014).

2. K. Katoh, D. M. Standley, MAFFT Multiple Sequence Alignment Software Version 7: Improvements in Performance and Usability. Mol. Biol. Evol. 30, 772–780 (2013).

3. N. C. Rockwell, S. S. Martin, J. C. Lagarias, Identification of Cyanobacteriochromes Detecting Far-Red Light. Biochemistry 55, 3907–3919 (2016).

4. N. C. Rockwell, S. S. Martin, J. C. Lagarias, Identification of DXCF cyanobacteriochrome lineages with predictable photocycles. Photochem. Photobiol. Sci. 14, 929–941 (2015).

5. J. Zhang, et al., Fused-Gene Approach to Photoswitchable and Fluorescent Biliproteins. Angew. Chem. Int. Ed. 49, 5456–5458 (2010).

6. Z. Otwinowski, W. Minor, “[20] Processing of X-ray diffraction data collected in oscillation mode” in Methods in Enzymology, Macromolecular Crystallography Part A., Jr. Charles W. Carter, Ed. (Academic Press, 1997), pp. 307–326.

7. E. S. Burgie, J. M. Walker, G. N. Phillips Jr., R. D. Vierstra, A Photo-Labile Thioether Linkage to Phycoviolobilin Provides the Foundation for the Blue/Green Photocycles in DXCF-Cyanobacteriochromes. Structure 21, 88–97 (2013).

8. P. D. Adams, et al., PHENIX: a comprehensive Python-based system for macromolecular structure solution. Acta Crystallogr. D Biol. Crystallogr. 66, 213–221 (2010).

9. P. Emsley, K. Cowtan, Coot: model-building tools for molecular graphics. Acta Crystallogr. D Biol. Crystallogr. 60, 2126–2132 (2004).

10. W. Humphrey, A. Dalke, K. Schulten, VMD: Visual Molecular Dynamics. J Molec Graph. 14, 33–38 (1996).

11. W. L. DeLano, Pymol: An open-source molecular graphics tool. CCP4 Newsletter On Protein Crystallography. 82–92 (2002).

